# Predicting woodland bird species habitat with multi-temporal and multisensor remote sensing data

**DOI:** 10.1101/2024.11.28.625964

**Authors:** Rachel J Kuzmich, Ross A Hill, Shelley A Hinsley, Paul E Bellamy, Ailidh E Barnes, Markus Melin, Paul M Treitz

## Abstract

Remote sensing data capture ecologically important information that can be used to characterize, model and predict bird habitat. This study implements fusion techniques using Random Forests (RF) with spectral Landsat data and structural airborne laser scanning (ALS) data to scale habitat attributes through time and to characterize habitat for four bird species in dynamic young forest environments in the United Kingdom. We use multi-temporal (2000, 2005, 2012/13, 2015) multi-sensor (Landsat and ALS) data to (i) predict structural attributes via pixel-level fusion at 30 metre spatial resolution, (ii) model bird habitat via object-level fusion and compare with models based on ALS, Landsat and predicted structural attributes, and (iii) predict bird habitat through time (i.e., predict 2015 habitat based on 2000-2012 data). First, we found that models predicting mean height from spectral information had the highest accuracy, whilst maximum height, standard deviation of heights, foliage height diversity, canopy cover and canopy relief ratio had good accuracy, and entropy had low accuracy. The green band and the normalized burn ratio (NBR) were consistently important for prediction, with the red and shortwave infrared (SWIR) 1 bands also important. For all structural variables, high values were underpredicted and low values were overpredicted. Second, for Blue Tit (*Cyanistes caeruleus*) and Chaffinch (*Fringilla coelebs*), the most accurate model employed Landsat data, while object-level fusion performed best for Chiffchaff (*Phylloscopus collybita*) and Willow Warbler (*Phylloscopus trochilus*). ALS mean, maximum and standard deviation of heights and Landsat tasseled cap transformations (TCT) (i.e., wetness, greenness and brightness) were ranked as important to all species across various models. Third, we used our models to predict presence in 2015 and implemented a spatial intersection approach to assess the predictive accuracy for each species. Blue Tit and Willow Warbler presences were well predicted with the Landsat, ALS, and objectlevel fusion models. Chaffinch and Chiffchaff presences were best predicted with the ALS model. Predictions based on pixel-level predicted structure surfaces had low accuracy but were acceptable for Chaffinch and Willow Warbler. This study is significant as it provides guidance for Landsat and ALS data application and fusion in habitat modelling. Our results highlight the need to use appropriate remote sensing data for each study species based on their ecology. Object-level data fusion improved habitat characterization for all species relative to ALS, but not to Landsat for Blue Tit and Chaffinch. Pixel-level fusion for predicting structural attributes in years where ALS data are note available is increasingly being used in modelling but may not adequately represent within-patch wildlife habitat. Finally, incorporating predicted surfaces generated through pixel-level fusion in our habitat models yielded low accuracy.

**Highlights:**

- We used object- and pixel-level fusion with ALS and Landsat to examine bird habitat
- Pixel-level fusion predicted surfaces yielded low accuracy in habitat models
- Best models: Landsat (Blue Tit, Chaffinch); fusion (Chiffchaff, Willow Warbler)
- Best prediction: ALS (Chaffinch, Chiffchaff)
- Best prediction: ALS, Landsat, object-level fusion (Blue Tit, Willow Warbler)

**Graphical abstract:** 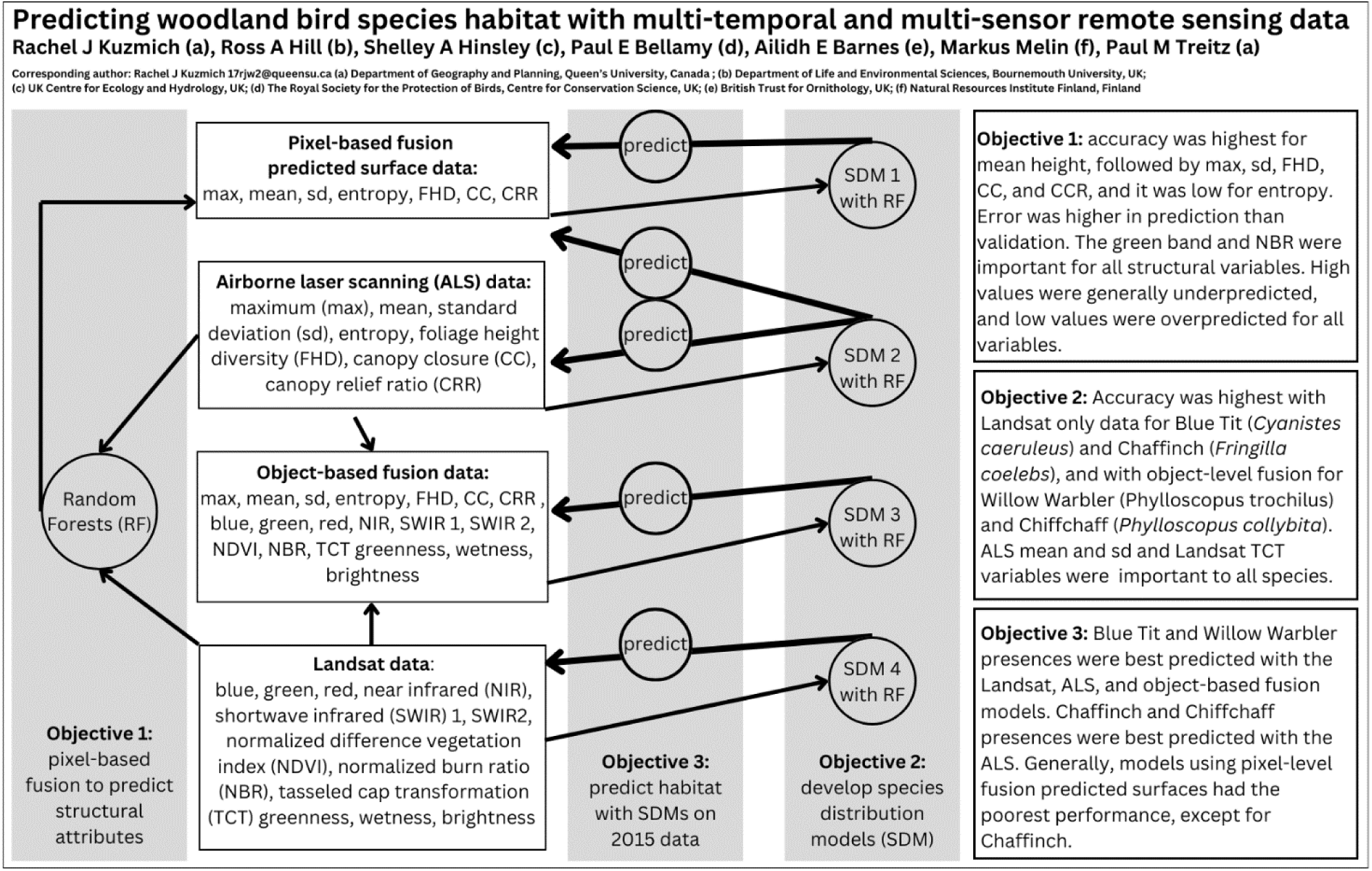

## 1 Introduction

Remote sensing data have become widely used in ecological applications due to their ability to capture ecologically important details at multiple spatiotemporal scales (Kerr & Ostrovsky, 2003; Wang et al., 2010). Field-based methods to collect environmental data in situ record highly detailed information but may experience logistical and financial constraints that limit collection to fine spatial and temporal extents (Wood et al., 2013) and potentially lead to biases (e.g., due to site selection based on accessibility Guillera-Arroita et al., 2015). Remote sensing, on the other hand, can capture ecologically meaningful information at multiple scales across broad spatial, and increasingly temporal, extents (Estes et al., 2018; Toth & Jóźków, 2016). The information and measurements captured by sensors make it possible to describe objects or surfaces (in 2D or 3D), over time.

For forest ecosystems, information derived from remote sensing data can characterize structure, composition and function (Balestra et al., 2024). For instance, the Normalized Difference Vegetation Index (NDVI) uses spectral reflectance information from the red and near infrared spectral bands to characterize vegetation (Huang et al., 2021; Rouse et al., 1974). This is ecologically important because NDVI values are positively correlated with biomass (Freitas et al., 2005; Zhu & Liu, 2015), productivity (Erasmi et al., 2021) and forest structure/complexity (Fiore et al., 2020). NDVI can also be related to plant species richness (Pau et al., 2012), and the occurrence, abundance or movement of wildlife species (Pettorelli et al., 2011; Regos et al., 2022). Spectral remote sensing data can also be applied to classify forest types (Adams et al., 2020; Dostálová et al., 2021), extract forest phenology information (Berra & Gaulton, 2021), assess forest ecotones (Hill et al., 2007; de Klerk et al., 2018), and characterize ecologically important aspects of horizontal vegetation structure (e.g., canopy cover; Guan et al., 2023). This is generally accomplished initially through object-oriented or pixel-based image classification using reflectance data, followed by extraction of ecologically meaningful variables. For instance, an object-based image analysis (OBIA) was used to assess post-war secondary forest regeneration, canopy cover change, and connectivity between ancient and regenerating woodlands in Poland using a combination of historic aerial photographs and satellite imagery (Jabs-Sobocińska et al., 2021). Forest ecosystems can also be characterized with 3D light detection and ranging (lidar) data to estimate measurements of canopy height (Mielcarek et al., 2018), describe the horizontal and vertical distribution of vegetation (Tang et al., 2019), derive foliar attributes (i.e., leaf area index; Richardson et al., 2009), and model the structural attributes of forest stands influencing wildlife species habitat (Martinuzzi et al., 2009; Hagar et al., 2020). Hereon, we will use the airborne laser scanning (ALS) convention when referring to lidar data collected using airborne platforms. Temporal characteristics can be described when data are available at multiple time steps, adding potentially a fourth dimension (4D) of information.

Remote sensing data are captured at a specific location and time, which enables a precise characterization of contemporary conditions, whereas time series can characterize the history of a place (Bolton et al., 2020), which is required to identify changes (e.g., disturbances; Huang et al., 2010) and to extract trends (e.g., changes to canopy cover; Vogeler et al., 2018). Overall, remote sensing offers data that can identify, characterize, quantify, model, and monitor ecologically important aspects of the environment that would be challenging to accomplish by other means.

The contribution of remote sensing to provide ecologically meaningful information related to the environmental characteristics of bird habitat was recognized decades ago (Nelson et al., 1971). Variables describing environmental characteristics of habitat have been extracted from remote sensing data and used to identify, characterize, monitor, and predict bird habitat (Gottschalk et al., 2005). These variables include land-use/land-cover classes (e.g., at the landscape scale; Fuller et al., 2005; although error propagation can be an issue; Cánibe et al., 2022), vegetation species composition (Buchanan et al., 2005), aspects of horizontal and vertical vegetation structure (Carrasco et al., 2019), topography (Kosicki, 2017), climate (Goetz et al., 2014), and oftentimes a combination thereof (e.g., edge effects, vegetation structure and composition; Broughton et al., 2012), all of which may influence bird species occurrence, abundance and distribution. While Landsat data have been used to examine aspects of bird habitat for decades (e.g., Palmeirim, 1988), lidar data are being used more frequently due to increasing availability (Bakx et al., 2019), though the use of structural attributes in bird habitat modelling pre-dates the advent of lidar technology (Dunlavy, 1935; Lack, 1933; MacArthur & MacArthur, 1961; MacArthur et al., 1962; van Dorp & Opdam, 1987). When coupled with wildlife species location information, remote sensing data can be used to develop a species distribution model (SDM). Indeed, SDMs increasingly incorporate remote sensing data to model the relationships between species and the environmental characteristics of their habitat (e.g., a species’ realized niche; Koma et al., 2021), often with the goal of predicting the occurrence of suitable habitat across space and/or time (i.e., to support conservation actions; Arenas-Castro & Sillero, 2021).

As suggested above, the data collected from different remote sensing systems (sensors and platforms) contain different information and measurements. Fusion is a broadly used term, beyond its use in remote sensing, referring to the implementation of methods of data combination, synergy, registration, integration, and calibration/validation (Balestra et al., 2024). Multi-sensor fusion has been used to estimate characteristics of surfaces, objects or changes (Huang et al., 2020), to improve classification accuracy (Guan et al., 2017), and to address information gaps in spectral, spatial, and temporal dimensions (or some combination thereof; e.g., see reviews on spatiotemporal fusion in Belgiu & Stein, 2019; Coops et al., 2021). While consensus on a precise definition of remote sensing data fusion is lacking and different studies have used this term to mean different things (Balestra et al., 2024), the goal is generally to estimate direct measurements or derived attributes from one sensor using information from another sensor, or to combine the information from multiple sensors to better characterize surfaces, objects or changes. In the application of habitat modelling, multi-sensor data fusion has been used to make predictions across space and time (Swatantran et al., 2012), to better characterize habitat and improve classification accuracy (Adams & Matthews, 2018; Vogeler et al., 2023).

Data fusion is generally done at the pixel-, feature-, or decision-level (Xiao et al., 2023) although other levels have been defined in specific applications. For instance, category-level fusion was used to distinguish foreground and background features with similar spectral characteristics (Zheng et al., 2022). A recent review of lidar fusion to estimate forest attributes includes datalevel and feature-level fusion, the latter of which is further subdivided into pixel-based and object-based fusion (Balestra et al., 2024). A further review that focused on fusion algorithms included signal-level fusion, which is akin to data-level fusion, but specifies the type of data (Dong et al., 2009). Remote sensing data fusion has been implemented using various approaches including spatial methods (i.e., k-nearest neighbor; Ahmed et al., 2016), statistical models (i.e., regression; Popescu & Wynne, 2004), machine learning (i.e., Random Forests; Veerabhadraswamy et al., 2021), and deep learning (i.e., convolutional neural networks; Chen et al., 2017).

By leveraging the relationship between structural ALS variables and spectral Landsat variables, it is possible to scale variables across space and time. This type of upscaling may be useful to predict forest structural characteristics in years when ALS data are not available. Typically, ALS data provide 3D structural information at a high spatial but low temporal resolution, whereas Landsat data provide spectral information at a lower spatial resolution but much greater temporal frequency, assuming cloud cover is not a persistent issue. Spaceborne lidar data from the Global Ecosystem Dynamics Investigation (GEDI) instrument have been integrated with spaceborne spectral data to scale lidar measurements (i.e., fusion with Landsat or Sentinel to estimate forest height; Potapov et al., 2021; Wang et al., 2024; or biomass; Tamiminia et al., 2024), and combined with multiple data sources to model habitat and biodiversity (i.e., Elliott et al., 2024).

Fusion of ALS and spectral data has been used to enhance the characterization and classification of habitat. As discussed above, various aspects of forest ecosystems can be described with different types of remote sensing data, all of which may be important in explaining wildlife species’ occurrence, abundance and distribution. Multi-sensor fusion leverages the complementary information of individual sensors (Swatantran et al., 2012), which has been found to increase classification accuracy. For instance, a Landsat-ALS fusion model yielded a classification accuracy improvement of 10 percentage points over Landsat alone, and even higher accuracy gains for the shrubland class which was important to shrubland-dependent bird species (Adams & Matthews, 2018). Similarly, the fusion of within-patch ALS variables, landscape-scale Landsat variables, and a radar-derived forest degradation index improved SDM accuracy for multiple bird species of conservation interest (Singh et al., 2017). Furthermore, Landsat-derived NDVI, and ALS-derived horizontal, vertical, and topographic variables were all found to contribute to bird models, although the relative importance varied across study sites and nesting guilds (Vogeler et al., 2014). Finally, remote sensing data fusion has improved habitat classification and characterization accuracy for birds in other ecosystems (e.g., tundra; Boelman et al., 2016).

Despite the advances in fusion techniques and applications, some challenges remain. Some of these challenges are practical, relating to radiometric calibration, atmospheric correction, data alignment and harmonization. Much of the pre-processing of remote sensing data that are to be used in data fusion addresses these practical challenges (Dwyer et al., 2018). Other challenges relate to specific environments, for instance, ecologically important aspects of young forests remain challenging to characterize due to their heterogeneous structure and mixed vegetation species composition (Ahmed et al., 2016), while others are application specific (e.g., for SDM development and interpretation, as discussed below).

For meaningful interpretation of any SDM that includes remote sensing data fusion, there is a dual challenge to represent both ecologically important aspects of the environment (Koma et al., 2022; Mod et al., 2016) and characteristics that are important to the ecology of the study species (Austin, 2002; Elith & Leathwick, 2009; MacNally, 2000). Furthermore, patterns of species distribution are the result of multiple factors impacting individual occurrence across multiple spatiotemporal scales (Fournier et al., 2017; Hallman & Robinson, 2020; Stevens & Conway, 2019). The availability and applicability of remote sensing data at the corresponding relevant scales is uncertain (Skidmore et al., 2021). Due to the spatial resolution of Landsat data, the application in finer spatial extents is less clear, with such data suggested for regional or landscape scale habitat studies (Bergen et al., 2007). In other words, Landsat pixels may be larger than some finer environmental characteristics and the value of each pixel will represent multiple features within that pixel (Cracknell, 1998).

The purpose of this study is to examine the accuracy of bird habitat modelling using multitemporal multi-sensor remote sensing data fusion techniques at two study sites undergoing woodland succession in Cambridgeshire, United Kingdom (UK). The study sites were selected because they are relatively young forests that are dynamic across time. Due to their complex and heterogenous structure, young forest stands are challenging to model with multi-sensor remote sensing data (Ahmed et al., 2015) but represent critical habitat for many bird species (Adams & Matthews, 2018). It is therefore essential to improve our understanding of the contribution of remote sensing fusion techniques to model and predict bird habitat in these environments. Blue Tit (*Cyanistes caeruleus*), Chaffinch (*Fringilla coelebs*), Chiffchaff (*Phylloscopus collybita*), and Willow Warbler (*Phylloscopus trochilus*) were selected, given that they occur frequently and provide sufficient data for analysis, while also representing species with different habitat associations. We use a time series of Landsat and ALS data to test two fusion methods that can be implemented in habitat modelling: (i) pixel-level fusion applied to predict structural attributes from spectral data; and (ii) object-level (i.e., presence and pseudo-absence sampling plots) fusion applied to model and predict breeding season habitat using SDMs developed with ALS, Landsat, fused, and the predicted structural attribute data.

Our specific objectives are to: 1) assess the ability of Landsat spectral data to predict ALS structural characteristics; 2) compare the capacity for Landsat-derived spectral data SDMs, ALSderived structural data SDMs, and structural-spectral fusion SDMs to characterize the breeding season habitat of each study species; and 3) compare the predictive accuracies of SDMs with structural (i.e., ALS), spectral (i.e., Landsat), fused (i.e., ALS and Landsat) and predicted (i.e., Landsat predicted structural attributes) data. Multi-temporal multi-sensor remote sensing data fusion has become widespread in land use/land cover applications (e.g., see reviews on the stateof-the-art in land use/land cover with remote sensing by Pandey et al., 2021), and represents an important area of further research for habitat modelling and prediction (Acebes et al., 2021). This study is significant as it will provide guidance for Landsat and ALS data use and fusion in habitat modelling, notably for bird species in dynamic young forest environments.

## 2 Materials and methods

### 2.1 Study area

This study focused on two study sites, ‘New Wilderness’ (2.1 ha) and ‘Old Wilderness’ (3.9 ha) (Boughton et al., 2021), in Cambridgeshire, UK (52° 24’N, 0° 14’W) (Figure 1). These sites are approximately 600 metres apart and are adjacent to Monks Wood National Nature Reserve, an ancient remnant woodland. The dominant land use in the broader landscape is agricultural with some small ancient woodland fragments and human infrastructure elements.

**Figure 1.**
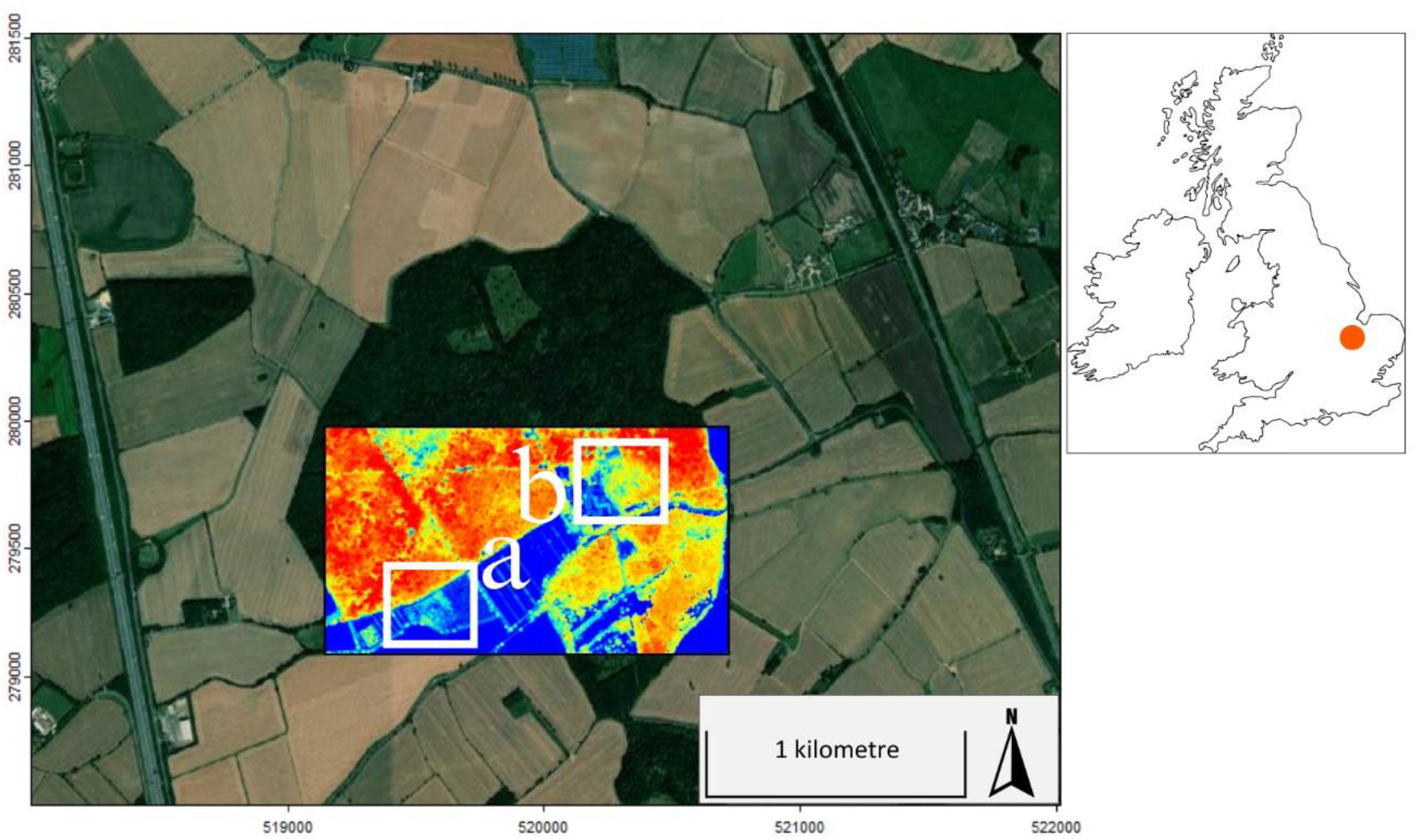
Study area map with the 2015 airborne laser scanning (ALS) data over the region of interest encompassing the: a) New Wilderness and b) Old Wilderness study sites.

Both sites were actively managed agricultural fields that have been abandoned and had minimal intervention, other than for path clearance. The Old Wilderness was abandoned in 1961 and the New Wilderness in 1996, both undergoing passive rewilding (Broughton et al., 2021). These sites were selected for analysis as they are dynamic across time. Woody vegetation density, cover, and height all increased as these sites were rapidly colonized, initially by shrub species and later with tree species (Broughton et al., 2021). Hawthorn (*Crataegus spp*.) and bramble (*Rubus fruticosus*) are common shrub species and Pedunculate Oak (*Quercus robur*) represents more than fifty percent of live trees in both wilderness sites (Broughton et al., 2021). Other common tree species in the wilderness sites include Elm (*Ulmus spp*.), Common Ash (*Fraxinus excelsior*) and Silver Birch (*Betula pendula*).

### 2.2 Remote sensing data and preprocessing

This research uses a time series of ALS and Landsat data. A summary of acquisition details is included in Table 1 for all remote sensing data.

**Table 1.** Overview of airborne laser scanning (ALS) and Landsat data acquisitions: Optech Airborne Laser Terrain Mapper (ALTM) and Leica Airborne Laser Scanner, Landsat 5 Thematic Mapper (TM) and Landsat 8 Operational Land Imager (OLI).

| Source | Sensor | Date | Spatial resolution |
| --- | --- | --- | --- |
| ALS | Optech ALTM 1210 | 2000-06-10 | 2 returns, 0.27 points/m<sup>2</sup> |
|  | Optech ALTM 3033 | 2005-06-26 | 2 returns, 0.45 points/m<sup>2</sup> |
|  | Leica ALS50-II | 2012-09-15 | 4 returns, 10.54 points/m<sup>2</sup> |
|  | Leica ALS50-II | 2015-06-22 | 4 returns, 2.1 points/m<sup>2</sup> |
| Landsat | TM | 2000-06-18 | 30 m |
|  | TM | 2005-06-09 | 30 m |
|  | OLI | 2013-07-17 | 30 m |
|  | OLI | 2015-08-08 | 30 m |

#### 2.2.1 ALS data acquisitions and preprocessing

ALS data were collected in 2000, 2005, 2012 and 2015 during the leaf-on period (Table 1). Different ALS discrete return systems with different configurations were used; the details of which have been described elsewhere (see Hill & Hinsley, 2015; Kuzmich et al., 2021). While differences in the number of returns and point density (noted in Table 1) mean that later acquisitions contain greater detail, notably the 2012 ALS point cloud, variables extracted from the ALS data are not significantly impacted (Lim et al., 2008; Treitz et al., 2012).

Pre-processing of ALS data was completed using the lidr package in R (Roussel & Auty, 2024). As the 3D point cloud data covered a broader geographic area, all data were clipped to a region of interest that contained both study sites with a buffer to facilitate processing (shown in Figure 1). All point clouds contained some duplicated points which were removed. The point clouds were then separated into ground and non-ground points for each acquisition using the cloth simulation function (Zhang et al., 2016). Subsequently, a digital terrain model (DTM) was generated from the ground points via kriging. A DTM with a 1 m resolution was generated for each of the ALS acquisitions, and pixel values were interpolated using the ten nearest neighbouring points. To confirm accuracy, the classified ground points and the generated DTM were visually inspected and compared to a terrain model available from the Centre for Environmental Data Analysis, UK. The ALS data for each year were normalized by subtracting the point clouds from the generated DTM. Outliers were removed using a progressive filtering technique to omit ALS points that were located high above the canopy or those with negative values below the DTM. This process resulted in four terrain normalized point clouds over the study sites, one for each ALS acquisition, where the value of each point represents height above ground level (agl).

#### 2.2.2 Landsat data acquisitions and preprocessing

Four Landsat images were obtained between 2000 and 2015. These data were collected during peak vegetation and visually inspected to confirm the absence of cloud cover or shadows over the study sites. Landsat data were selected to match the ALS acquisition dates as closely as possible; however, if a corresponding image was not available due to cloud contamination (i.e., 2012), we substituted a suitable image acquired the following year (i.e., 2013).

All Landsat images are Collection 2 Level-2 (C2L2) science products from the United States Geological Survey (USGS) and were acquired via EarthExplorer. The 2000 and 2005 data were collected with Landsat 5 Thematic Mapper (TM) and 2013 and 2015 with Landsat 8 Operational Land Imager (OLI) (Table 2). Despite subtle differences across Landsat sensors, they collect complementary information (Mancino et al., 2020) and the C2L2 images are intended for use in time series (Crawford et al., 2023). Noted that we used only the shared bands from TM and OLI, the coastal aerosol, panchromatic and cirrus bands from OLI were not used.

**Table 2.**
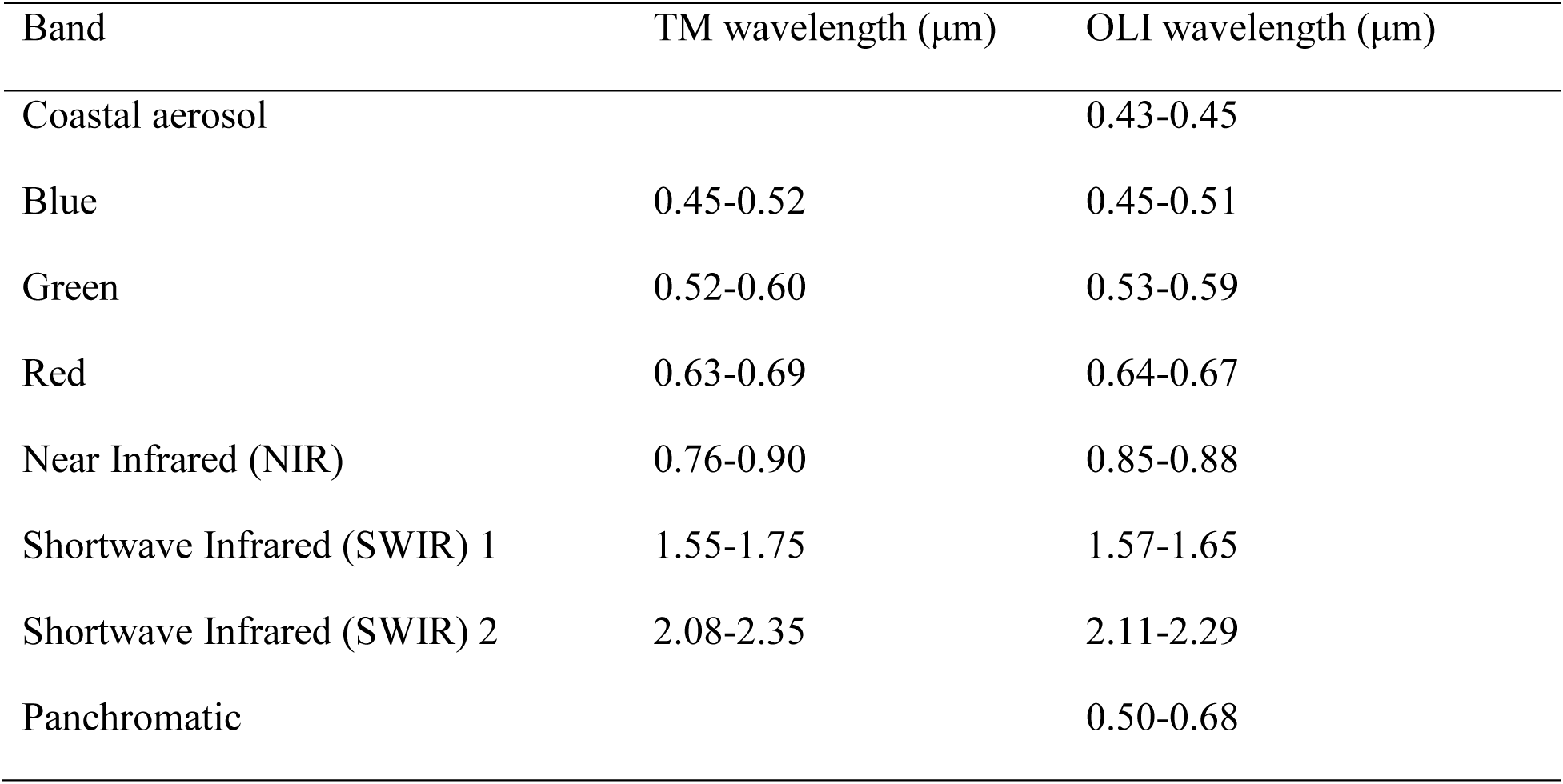
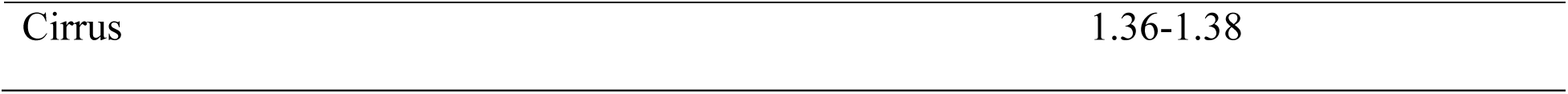
Spectral information of Landsat 5 Thematic Mapper (TM), and Landsat 8 Operational Land Imager (OLI) reflectance data.

Following the procedure outlined in Young et al. (2017), the digital numbers of all Landsat data were converted to at-sensor radiance and subsequently to top-of-atmosphere (TOA) reflectance. Given that this analysis used multi-temporal multi-sensor data (e.g., TM and OLI), the conversion to radiance makes data comparable. The correction to TOA accounts for power, distance and elevation angle of the sun which varies with time and by location. These preprocessing steps were completed manually in R following Goslee (2011) with calibration coefficients (e.g., mean exoatmospheric solar irradiance) provided in Chander et al. (2009) and Januar et al. (2020). To improve the temporal continuity between Landsat acquisitions, we developed per band functions for relative calibration with ordinary least squares regression (Roy et al., 2016; Young et al., 2017). The coordinate reference system for all Landsat data was verified and reprojected using the sf (Pebesma & Bivand, 2023) and terra (Hijmans, 2024) packages to ensure consistency across all data. Landsat data were clipped to the same region of interest as the ALS data encompassing both study sites (corresponding to the ALS overlay in Figure 1).

### 2.3 Bird data and preprocessing

The bird data comprised a series of digitized observations of all individuals (either seen or heard) collected across multiple surveys throughout the spring to early summer bird breeding season at the Old and New Wilderness sites. Surveys were conducted by trained experts when weather conditions were favourable using a spot mapping method based on the Common Birds Census (CBC) of the British Trust for Ornithology (Marchant, 1983). There are gaps in the bird data either due to lack of surveying (e.g., New Wilderness surveying began in 2001) or because of lack of detection (e.g., no Willow Warblers were observed in 2012 at Old Wilderness).

If the same individual for any of the four bird species studied was observed multiple times during the same survey, only the first observation was retained. To avoid overlapping bird habitat sampling plots, bird observations were removed (selected randomly) to ensure a minimum distance of 30 metres between observations. A balanced number of background points representing pseudo-absence locations were generated for each bird species using the same minimum distance requirement to avoid overlap.

This study used Blue Tit, Chaffinch, Chiffchaff and Willow Warbler data from 2000-2002, 2005-2007, 2012-2014, and 2015-2017. These dates correspond to the 2000, 2005, 2012/2013 and 2015 ALS and Landsat data acquisitions. We used bird data from the same year as the remote sensing acquisitions and the two subsequent years because birds may exhibit site fidelity or a delayed response to structural changes (Bellamy et al., 2000; Jakobsson, 1988; Schlossberg, 2009). We selected these bird species to represent a range of habitat preferences. As a cavity nester, the Blue Tit is a woodland generalist typical of more mature sites, while the Chaffinch occurs widely across woodland, scrub and farmland. The Willow Warbler (in lowland England) is typical of early successional and edge habitat, avoiding well-grown trees and closed canopy mature woodland while the Chiffchaff commonly occurs in woodland with a good shrub/herb layer component and shrubby/mature hedgerow habitats with trees.

### 2.4 Fusion to predict structural attributes from spectral data

Our first objective is to predict ALS structural characteristics from Landsat spectral data (both individual band information and computed metrics). This type of fusion was done at the pixel level to generate a fused image (e.g., raster surface) with estimated pixel values (Dong et al., 2009). We randomly split the model development data to reserve 30% for validation. We also used temporal cross-validation (Roberts et al., 2017) to test the predictive ability of our model. Note that the predicted structural surfaces will also be used to assess the predictive ability of SDMs developed with ALS data to support our second and third objectives. All variables used in this study are described in Table 3.

**Table 3.**
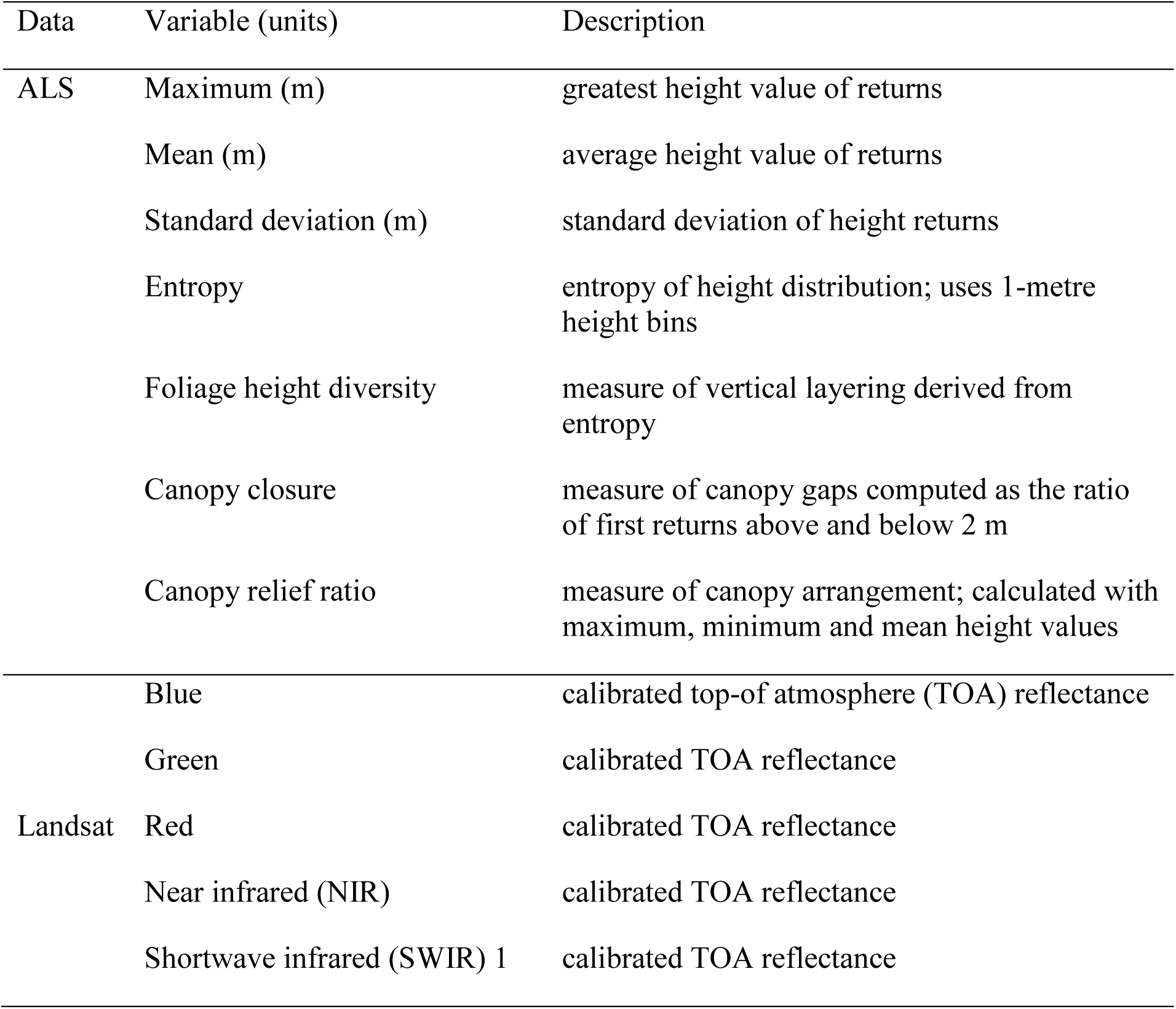
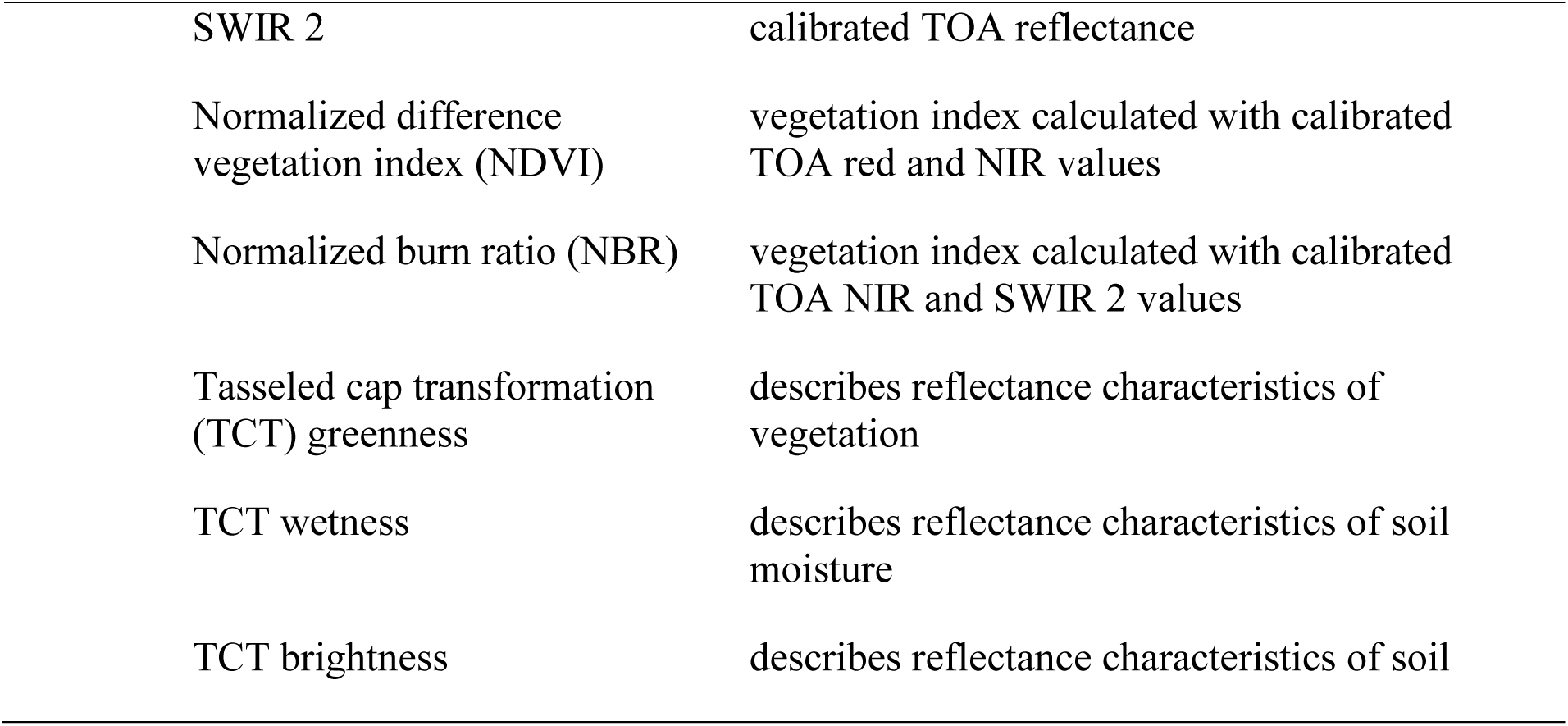
Summary of variables extracted from the airborne laser scanning (ALS), Landsat 5 Thematic Mapper (TM) and Landsat 8 Operational Land Imager (OLI) data.

| Data | Variable (units) | Description |
| --- | --- | --- |
| ALS | Maximum (m) | greatest height value of returns |
|  | Mean (m) | average height value of returns |
|  | Standard deviation (m) | standard deviation of height returns |
|  | Entropy | entropy of height distribution; uses 1-metre height bins |
|  | Foliage height diversity | measure of vertical layering derived from entropy |
|  | Canopy closure | measure of canopy gaps computed as the ratio of first returns above and below 2 m |
|  | Canopy relief ratio | measure of canopy arrangement; calculated with maximum, minimum and mean height values |
| Landsat | Blue | calibrated top-of atmosphere (TOA) reflectance |
|  | Green | calibrated TOA reflectance |
|  | Red | calibrated TOA reflectance |
|  | Near infrared (NIR) | calibrated TOA reflectance |
|  | Shortwave infrared (SWIR) 1 | calibrated TOA reflectance |

|  |  |
| --- | --- |
| SWIR 2 | calibrated TOA reflectance |
| Normalized difference vegetation index (NDVI) | vegetation index calculated with calibrated TOA red and NIR values |
| Normalized burn ratio (NBR) | vegetation index calculated with calibrated TOA NIR and SWIR 2 values |
| Tasseled cap transformation (TCT) greenness | describes reflectance characteristics of vegetation |
| TCT wetness | describes reflectance characteristics of soil moisture |
| TCT brightness | describes reflectance characteristics of soil |

#### 2.4.1 Rasterize ALS data

ALS data were transformed from a 3D point cloud to multiple raster surfaces at a 30 metre resolution, each representing one of the structural characteristics described in Table 3. This was accomplished in R using the lidr package (Roussel & Auty, 2024). Data were projected to the same coordinate reference system as the Landsat data and clipped to a smaller region of interest to remove any potential edge effects (e.g., from DTM generation). As a result, the spatial resolution and alignment of the generated ALS raster surfaces was consistent and matched the corresponding Landsat data.

Maximum and mean values summarize the height of ALS returns within pixels to describe the vertical distribution of vegetation. The lidr package includes a function to compute entropy (Roussel et al., 2020; Shannon, 1948), and foliage height diversity (FHD) was calculated by multiplying entropy by the log of the maximum ALS height return value (z_max_) (Atkins et al., 2023; MacArthur & MacArthur, 1961; MacArthur et al., 1962).

Along with the standard deviation of heights, these variables describe heterogeneity in the vertical distribution of ALS points, with entropy and FHD specifically representing the distribution of layers of vegetation in the forest. Canopy closure (CC) describes the horizontal distribution of ALS returns using a threshold to separate vertical vegetation layers (Ahmed et al., 2015) and represents the proportion of lower (i.e., understorey) vegetation that is obscured by the upper canopy (Vogeler et al., 2018). It was calculated as follows:

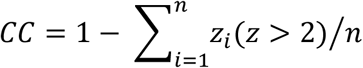

where n refers to the number of returns and z refers to the ALS height values. Note that here CC was calculated using the ratio of ALS returns above and below 2 m. This threshold was selected to represent the vegetation structure at the two study sites and to separate the upper canopy (e.g., overstorey) from the lower (e.g., understorey) vegetation.

The canopy relief ratio (CRR), the relative arrangement of vegetation within the bird habitat sampling plots (Atkins et al., 2023; Hawryło et al., 2017), was calculated as follows:

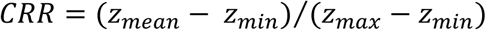

where z refers to the height of the ALS return, and mean, min and max refer respectively to the mean, minimum and maximum height values for the ALS returns.

#### 2.4.2 Model development

We used Random Forests (RF) as a regression approach to model the relationship between Landsat spectral data and rasterized ALS structural surfaces to make predictions. This method was selected because RF has previously been used to predict structural forest attributes with spectral data (e.g., Ahmed et al., 2015; Bolton et al., 2020). RF is a non-parametric machine learning classification and regression algorithm that develops multiple uncorrelated decision trees forming a forest (Breiman, 2001). Each decision tree is fitted with a bootstrap sample of the training data and grown through recursive random splits of the response data on predictor variables at each node (Biau & Scornet, 2016; Valavi et al., 2021). These splits are made to maximize within-node similarity and between-node dissimilarity so that each split increases purity. This algorithm makes no assumptions about the relationship between predictor and response variables (i.e., non-parametric), inherently includes interactions between variables (Cutler et al., 2007; Elith et al., 2008), and is insensitive to overfitting relative to other modelling methods (Belgiu & Drăguţ, 2016).

For each ALS structural variable (Table 3), we developed a RF model using three years of data (e.g., 2000, 2005, 2012). Model development data were randomly split to reserve 30% for validation. Data from one year was withheld from each model to be used for prediction of that year (e.g., 2015). This procedure produced four modelling scenarios, described in Table 4. Models were developed in R using the default tuning parameters with randomForest (Liaw & Wiener, 2002).

**Table 4.**
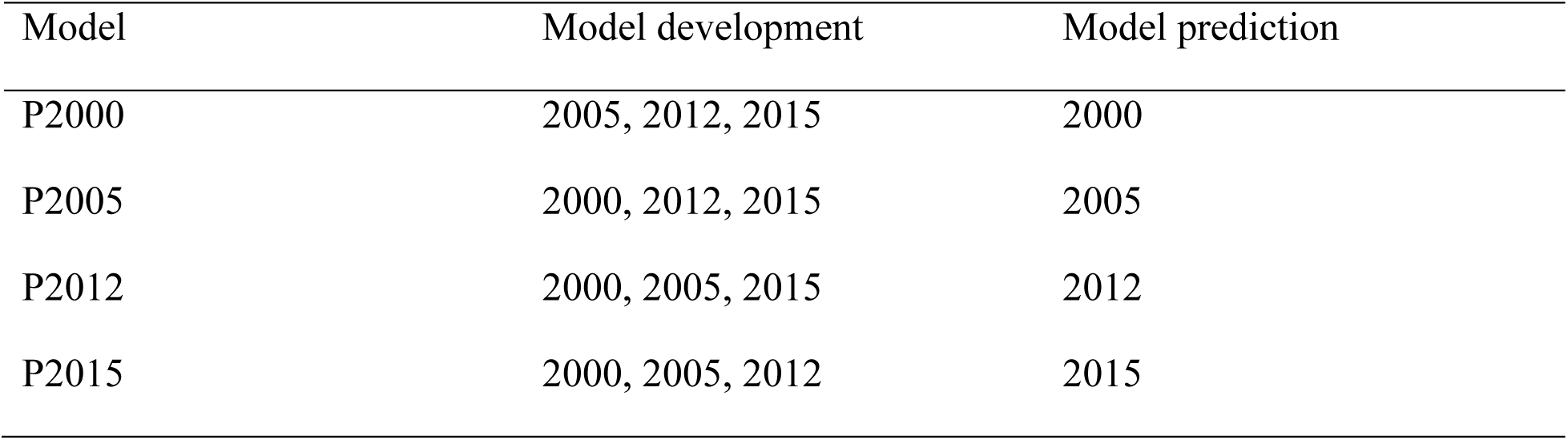
Modelling scenarios used in this study for prediction of structural attributes from spectral data, including the data years used for model development and for model prediction.

| Model | Model development | Model prediction |
| --- | --- | --- |
| P2000 | 2005, 2012, 2015 | 2000 |
| P2005 | 2000, 2012, 2015 | 2005 |
| P2012 | 2000, 2005, 2015 | 2012 |
| P2015 | 2000, 2005, 2012 | 2015 |

#### 2.4.3 Model assessment

For RF regression models, the output includes a measure of variation explained by the model and a percent increase in mean squared error for each variable in the model which can be used to identify important variables (Liaw & Wiener, 2002). Validation data were extracted from the same years of data that were used in development (70:30 split). We used the validation datasets to calculate root mean square error (RMSE) to assess the difference between the predicted and actual values. As our primary interest here is to predict structural attributes through time with spectral data and derived variables, RMSE was also calculated to assess how generalizable the developed models were for making predictions through time using temporal cross validation. Here RMSE represents the average absolute difference between the predicted structural attributes and the actual values from the ALS data. To understand the magnitude of the RMSE values, we also calculated relative RMSE (Bolton et al., 2020) which standardizes the absolute RMSE values to the range of actual values. Finally, to better understand the error associated with temporal prediction, we produced a series of ‘difference’ maps using the rasterized ALS data which were not included in model development and the generated predictive surfaces.

### 2.5 Fusion to model bird habitat with multi-sensor remote sensing data

Our second objective was to develop SDMs to model the breeding season habitat of each study species using Landsat data, ALS data, multi-sensor fusion, and generated predicted surfaces (from our first objective). Here the feature that we are characterizing is habitat using the sampling plots as objects. OBIA facilitates the fusion of multi-sensor data for improved classification relative to per-pixel analyses (Blaschke, 2010). While OBIA is widely used to classify features with clearly defined boundaries (Hossain & Chen, 2019), it can be used as an analytical tool to classify land cover (Evans & Costa, 2013; Jabs-Sobocińska et al., 2021; Onojeghuo & Onojeghuo, 2017), and we extend this to conceptualize bird species’ habitat (Glad et al., 2020). A key difference here is that, rather than using remote sensing spectral or structural similarity for image segmentation, objects were created using presence and pseudo-absence locations. We use object-level data fusion to extract and subsequently combine spectral and structural information, characterizing these objects from the remote sensing data (Xiao et al., 2023). In other words, ALS and Landsat candidate variables describing Blue Tit, Chaffinch, Chiffchaff and Willow Warbler breeding season habitat were extracted from the presence and pseudo-absence sampling plot objects and subsequently integrated in a fused SDM. To compare accuracy and assess any relative differences, we also developed SDMs for each bird species with data from Landsat and ALS separately, and with the generated predicted surfaces, resulting in four SDMs for each of the study species.

#### 2.5.1 Remote sensing variable extraction

All remote sensing variables were extracted using a 15-metre radius circular plot centred on bird presence and pseudo-absence locations (Hinsley et al., 2009; Kuzmich et al., forthcoming). The variables that were extracted and computed from the ALS and Landsat data are the same as those described in Table 3. Note that variables were calculated for the presence and pseudo-absence sampling plots (rather than being transformed to raster pixels). ALS sampling data were extracted from the normalized 3D point clouds using the lidr package in R (Roussel & Auty, 2024). For the Landsat data and derived variables (e.g., NDVI), sampling from the raster used an area-weighted mean to extract sampling plot values with the terra package in R (Hijmans, 2024). Some of the bird locations recorded by the surveys were in the central area of the study sites while others were at the edge. This meant that some of the sampling plots include greater variation in spectral attributes (as was the case with the structural attributes extracted from ALS).

From the Landsat data, we used individual band information and computed NDVI (Rouse et al., 1974), normalized burn ratio (NBR) (Key & Benson, 2006), and tasseled cap transformation (TCT) brightness, greenness and wetness (Crist & Cicone, 1984; Crist & Kauth, 1986). TCT components were computed in R using the RStoolbox package (Leutner et al., 2024), whilst NDVI and NBR were calculated manually in R.

NDVI was calculated as follows:

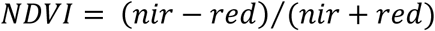

where nir and red refer to the corresponding NIR and red calibrated TOA reflectance values (e.g., respectively from Landsat 5 bands 3 and 4, and Landsat 8 bands 4 and 5).

NBR was calculated as follows:

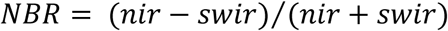

where nir and swir refer to the corresponding NIR and SWIR 2 calibrated TOA reflectance values (e.g., respectively from Landsat 5 bands 4 and 7, and Landsat 8 bands 5 and 7).

Individual band information was used as these have been found to be important for modeling bird habitat across multiple vegetation types and because this retains all information contained in the Landsat data (Adams & Matthews, 2018). NDVI was computed because it has been widely recognized for its ecological applications, is simple to compute, and has been found to be significant in habitat modelling (e.g., for bird diversity in Bonthoux et al., 2018; see also reviews by Leyequien et al., 2007; Pettorelli et al., 2011). While not widely used in bird habitat studies, the NBR has been used to characterize vegetation regeneration due to its association with vegetation structure (White et al., 2017). TCT brightness, greenness and wetness variables describe the spectral characteristics of soil, vegetation, and soil/vegetation moisture (Crist & Cicone, 1984; Crist & Kauth, 1986) and have been used to model characteristics of forests (Healey et al., 2005; Stankova & Avetisyan, 2024), aspects of bird habitat (Alessandrini et al., 2022; Helmer et al., 2010; Moreira et al., 2022), and to predict ALS-derived structural attributes (e.g., Bolton et al., 2020).

#### 2.5.2 Model development

We used RF to identify the remote sensing variables that were important for characterizing the breeding season habitat of Blue Tit, Chaffinch, Chiffchaff and Willow Warbler. Four separate models were developed for each bird species, one with Landsat variables, one with ALS variables, one with structural variables from the generated predicted surfaces, and a multi-sensor fusion model that combined ALS and Landsat data at the object-level. Here we are using RF to classify the sampling plot object data as either categorical presence or pseudo-absence variables. All models were developed with the 2000, 2005, and 2012/13 data. Data were randomly split so that 70% were used for training and 30% were used for validation. We used the default tuning parameters for the number of trees and the number of variables tried at each split. RF has become a common method in SDM development due to its ability to accurately model complex relationships between species and the environmental characteristics of their habitat derived from remote sensing data (e.g., recently for SDM development for bird species in Gábor et al., 2024; Huettman et al., 2024).

#### 2.5.3 Model assessment

For RF classification models, data omitted from the bootstrap sample (i.e., not used as training data) were used to calculate out-of-bag (OOB) error. RF also provides measures of variable importance based on mean decrease Gini (MDG) and mean decrease accuracy (MDA), which may be used to rank or select candidate variables (Genuer et al., 2010; Nicodemus, 2011; Song et al., 2021; Wang et al., 2016). Important variables were identified using measures of importance based on MDA values for each variable, as it is robust to correlation within the predictor variable set relative to MDG (Nicodemus, 2011). Variable importance was plotted for each model to facilitate interpretation using the randomForest package in R (Liaw & Wiener, 2002). We also submitted a validation dataset (e.g., 30% of data from 2000, 2005 and 2012) to the RF models to assess performance.

### 2.6 Bird habitat prediction

Our final objective was to predict Blue Tit, Chiffchaff, Chaffinch and Willow Warbler presence based on the structural and spectral characteristics of their habitat. Data from 2000, 2005 and 2012/13 were used to develop these SDMs (as described above) and predictions were made on the 2015 data. We considered five prediction scenarios (Table 5) for each of the four study species.

**Table 5.** Species distribution model (SDM) prediction (P) scenarios used in this study, specifying the data used for model development and prediction.

| Model | SDM development | SDM prediction |
| --- | --- | --- |
| P_ALS | ALS data | ALS data |
| P_Landsat | Landsat data | Landsat data |
| P_Comb | Object-level fusion (objective 2) | Fused data |
| P_Pred | Predicted surfaces (objective 1) | Predicted surface (objective 1) |
| P_ALSandPred | ALS data | Predicted surface (objective 1) |

Our first two SDM prediction scenarios used the two remote sensing data types separately. In other words, the SDM developed with ALS data was used for prediction on the 2015 ALS data (P_ALS), and the same for Landsat (i.e., P_Landsat). The third scenario used the multi-sensor fusion SDM that combined ALS and Landsat data at the object-level (objective 2) for prediction on a fused raster that combined the ALS and Landsat data (i.e., P _Comb). To be able to make predictions for the P _Comb scenario, we created a fusion raster surface in R that combined the 2015 rasterized ALS data and 2015 Landsat data on a pixel-by-pixel basis. Next, we used the SDM developed with the Landsat predicted structural attributes (objective 1) for prediction on the generated predicted surface (i.e., P_Pred). Finally, we used the SDM developed with ALS data for prediction on the predicted surface (i.e., P_ALSandPred) as both the training data and the predicted surface represent the same structural attributes. In other words, the predictions should be the same as the results of the first scenario if the prediction of structural attributes (objective 1) is perfect. A binary raster surface (i.e., presence, pseudo-absence) was created for each prediction in R.

#### 2.6.1 Prediction assessment

The accuracy of each prediction surface was assessed using the corresponding bird data from 2015-2017 and the same 15-metre radius circular plots centred on presence locations. We used a spatial intersection approach where presence polygons were overlaid onto the binary raster surface in R using the terra package (Hijmans, 2024). Given that these polygons intersected with multiple pixels, some of which may be predicted as presence or as absence, the polygon was considered correct if it intersected with any predicted presence pixels. We then manually calculated the predicted presence error as the proportion of presence sampling plots that did not intersect with any predicted presence pixels in R.

## 3 Results

### 3.1 Predicting structural characteristics from spectral data

In this study, we modelled the relationship on a pixel-by-pixel basis between Landsat spectral data and rasterized ALS data to create a fused image representing predicted structural attributes for four modelling scenarios (i.e., P2000, P2005, P2012, P2015; see Table 4). The variation explained by the RF models was greatest for mean height (0.80-0.83). Models predicting maximum height, standard deviation of heights, FHD, CC and CRR had good accuracy (0.71-0.81). Entropy had the lowest accuracy of all the modelled structural variables (0.58-0.66). The P2005 scenario, which was developed with 2000, 2012 and 2015 data, had the greatest variation explained by the RF models based on OOB sampling. The variation explained by all RF models is summarized in Table 6.

**Table 6.** Summary of variation explained by RF models based on out-of-bag (OOB) sampling for the four prediction scenarios.

| Variable (units) | P2000 | P2005 | P2012 | P2015 |
| --- | --- | --- | --- | --- |
| Maximum (m) | 0.78 | 0.80 | 0.78 | 0.77 |
| Mean (m) | 0.80 | 0.83 | 0.80 | 0.80 |
| Standard deviation | 0.74 | 0.76 | 0.72 | 0.70 |
| Entropy | 0.66 | 0.63 | 0.58 | 0.58 |
| Foliage height diversity: FHD | 0.79 | 0.81 | 0.77 | 0.75 |
| Canopy closure: CC | 0.79 | 0.81 | 0.75 | 0.75 |
| Canopy relief ratio: CRR | 0.75 | 0.75 | 0.71 | 0.72 |

Model accuracy was assessed using the validation dataset, which represented 30% of the data from the same years used in model development that were excluded from the training process. RMSE values were generally low for all the modelled structural variables (Table 7). For example, there was a prediction error of 3.32-3.59 metres in the maximum height estimations across all RF modelling scenarios, representing a relative RMSE error of 14-16 %. Overall, the lowest relative RMSE were associated with the P2005 modelling scenario for all variables.

**Table 7.** Summary of absolute and relative (rel) root mean square error (RMSE) based the validation dataset for the four scenarios (i.e., P2000, P2005, P2012, P2015).

| Variable (units) | P2000 (rel) | P2005 (rel) | P2012 (rel) | P2015 (rel) |
| --- | --- | --- | --- | --- |
| Maximum (m) | 3.59 (0.16) | 3.32 (0.14) | 3.47 (0.15) | 3.47 (0.16) |
| Mean (m) | 2.54 (0.16) | 2.28 (0.14) | 2.69 (0.16) | 2.62 (0.16) |
| Standard deviation (m) | 1.12 (0.17) | 1.10 (0.17) | 1.11 (0.19) | 1.12 (0.17) |
| Entropy | 0.16 (0.18) | 0.16 (0.18) | 0.17 (0.20) | 0.17 (0.19) |
| Foliage height diversity: FHD | 0.44 (0.16) | 0.42 (0.15) | 0.46 (0.17) | 0.44 (0.16) |
| Canopy closure: CC | 0.17 (0.17) | 0.16 (0.17) | 0.17 (0.18) | 0.16 (0.17) |
| Canopy relief ratio: CRR | 0.11 (0.16) | 0.11 (0.15) | 0.12 (0.18) | 0.11 (0.16) |

We used the same measure, RMSE, to assess the predictive ability of all RF models. The data used for prediction were from the years that were entirely excluded from model development and used only here for temporal cross-validation to test the predictive ability and generalizability of the models. Compared to the validation dataset, RMSE values were higher for all variables (Table 8). For instance, the RMSE associated with maximum height was 3.88-4.94 metres, which is 0.56-1.35 metres greater than the RMSE validation error values, and as high as a doubling of relative RMSE values (i.e., for P2005). For prediction, the lowest relative RMSE values were for the P2000 (maximum, mean, CRR), P2012 (entropy, CC, CRR) and P2015 (standard deviation, entropy, FHD, CC, CRR) modelling scenarios. The P2005 modelling scenario had the highest relative and absolute RMSE values.

**Table 8.** Summary of absolute and relative (rel) root mean square error (RMSE) based on the prediction dataset for each of the four scenarios (i.e., P2000, P2005, P2012, P2015).

| Variable (units) | P2000 (rel) | P2005 (rel) | P2012 (rel) | P2015 (rel) |
| --- | --- | --- | --- | --- |
| Maximum (m) | 3.88 (0.20) | 4.94 (0.28) | 3.97 (0.18) | 4.63 (0.27) |
| Mean (m) | 2.70 (0.16) | 3.86 (0.27) | 2.55 (0.18) | 3.32 (0.23) |
| Standard deviation (m) | 1.27 (0.23) | 1.45 (0.30) | 1.47 (0.28) | 1.21 (0.20) |
| Entropy | 0.21 (0.30) | 0.25 (0.36) | 0.17 (0.22) | 0.15 (0.22) |
| Foliage height diversity: FHD | 0.52 (0.20) | 0.68 (0.33) | 0.48 (0.19) | 0.47 (0.18) |
| Canopy closure: CC | 0.21 (0.22) | 0.25 (0.31) | 0.18 (0.20) | 0.18 (0.20) |
| Canopy relief ratio: CRR | 0.14 (0.20) | 0.15 (0.25) | 0.13 (0.20) | 0.12 (0.20) |

Important variables were identified using the percent increase in mean squared error for each variable in each modelling scenario (Table 9). Information from the green band and NBR were consistently the most important variables for all structural attributes, except for entropy which had the lowest accuracy of all the modelled structural attributes. The SWIR 1, red and SWIR 2 bands were repeated among the top three variables for multiple structural attributes. The NIR band and TCT wetness were among the top five variables for maximum, mean and standard deviation of heights, and for mean height, CC and CCR, respectively. The blue band, NDVI, TCT brightness and greenness were generally among the least important. Note however, that even the least important variables contribute to the model.

**Table 9.** The average rank and range of percent increase in mean square error (in brackets) for each variable across all scenarios. While all variables contributed to the accuracy of the models, the top three for each of the modelled structural variables are in bold to facilitate interpretation.

|  | Maximum | Mean | Standard deviation | Entropy | FHD | CC | CRR |
| --- | --- | --- | --- | --- | --- | --- | --- |
| Blue | 10<br>(17.32-20.16%) | 11<br>(17.59-18.60%) | 10<br>(16.92-20.38%) | 10<br>(14.59-19.10%) | 10<br>(15.58-18.19%) | 11<br>(14.72-15.80%) | 10<br>(16.30-17.36%) |
| Green | **1**<br>**(29.59-36.36%)** | **1**<br>**(28.44-36.56%)** | **1**<br>**(32.70-41.26%)** | 4<br>(21.34-27.32%) | **1**<br>**(30.19-35.15%)** | **1**<br>**(29.25-35.31%)** | **1**<br>**(27.33-31.69%)** |
| Red | 5<br>(19.05-27.60%) | **2**<br>**(20.19-32.26%)** | 4<br>(25.66-30.51%) | **2**<br>**(16.84-30.02%)** | 4<br>(17.88-29.80%) | **3**<br>**(16.37-30.47%)** | 5<br>(16.92-29.03%) |
| NIR | 5<br>(21.17-26.61%) | 4<br>(25.86-28.29%) | 5<br>(23.36-30.47%) | 6<br>(19.22-22.30%) | 7<br>(19.64-22.60%) | 6<br>(19.58-22.22%) | 6<br>(18.23-27.01%) |
| SWIR 1 | **3**<br>**(25.16-34.37%)** | 7<br>(18.42-26.91%) | **3**<br>**(27.86-24.60%)** | **1**<br>**(25.75-32.00%)** | **3**<br>**(19.49-31.35%)** | 4<br>(20.99-25.14%) | **3**<br>**(22.47-27.16%)** |
| SWIR 2 | 4<br>(23.65-27.58%) | 9<br>(16.03-21.00%) | 5<br>(22.95-28.41%) | **2**<br>**(22.58-28.69%)** | 5<br>(19.07-26.79%) | 9<br>(16.86-17.53%) | 9<br>(15.38-18.81%) |
| NDVI | 9<br>(20.31-23.93%) | 8<br>(18.99-22.01%) | 8<br>(20.05-23.13%) | 7<br>(15.75-20.21%) | 8<br>(17.16-20.49%) | 7<br>(16.64-22.22%) | 10<br>(16.08-20.25%) |
| NBR | **1**<br>**(28.89-36.65%)** | **3**<br>**(25.21-29.69%)** | **2**<br>**(30-28-36.01%)** | 4<br>(18.19-28.63%) | **2**<br>**(24.63-31.73%)** | **2**<br>**(20.63-27.57%)** | **2**<br>**(24.64-27.37%)** |
| TCT greenness | 8<br>(19.63-24.62%) | 6<br>(20.86-24.98%) | 7<br>(19.51-26.12%) | 10<br>(16.24-18.30%) | 9<br>(16.27-19.92%) | 8<br>(17.43-18.63%) | 8<br>(16.18-22.36%) |
| TCT wetness | 7<br>(19.49-27.39%) | 5<br>(20.26-27.21%) | 8<br>(19.22-26.58%) | 7<br>(15.73-23.19%) | 6<br>(17.92-26.54%) | 4<br>(20.37-26.96%) | 4<br>(22.13-27.02%) |
| TCT brightness | 10<br>(17.60-18.26%) | 10<br>(18.54-20.44%) | 11<br>(18.77-20.27%) | 11<br>(12.13-17.14%) | 10<br>(14.69-19.85%) | 10<br>(14.79-19.68%) | 7<br>(16.11-20.78%) |

Note: acronyms include near infrared (NIR), shortwave infrared (SWIR), normalized difference vegetation index (NDVI), normalized burn ratio (NBR), tasseled cap transformation (TCT), foliage height diversity (FHD), canopy cover (CC), and canopy relief ratio (CRR).

To better understand the errors in prediction, we have produced a series of surfaces for each of the predicted structural variables at the Old Wilderness study site for the P2015 scenario (see supplementary materials: Figures S1 to S7). A similar general pattern emerged across the predictions for all of the structural variables. Generally, the greatest difference between the predicted and actual values of the structural attributes occurs in the mature forest (north-east), the area featuring bare ground and buildings (south-west), or with the occurrence of variable vegetation structure associated with open areas such as canopy gaps and hedgerow features (south-east). Lower prediction error occurs in the central area of this region of interest, except for standard deviation of heights, which is where the Old Wilderness study site is located. Furthermore, lower prediction error is generally associated with intermediate values for all structural attributes, whereas the mature forest area (e.g., which has higher maximum height values) is under-predicted and the bare ground (e.g., which has lower maximum height values) is over-predicted. Overall, the range of predicted values was narrower than the actual values.

### 3.2 Modeling bird habitat with multi-sensor remote sensing data

The OOB and validation error for RF models developed using Landsat variables are summarized in Table 10. Both the OOB model error and the validation dataset error were lowest for Willow Warbler, followed by Chaffinch and Blue Tit, whilst Chiffchaff had the highest error. Presence and pseudo-absence OOB errors were equal for Blue Tit and Chaffinch, whereas OOB presence error was greater than pseudo-absence error for Chiffchaff and Willow Warbler.

**Table 10.**
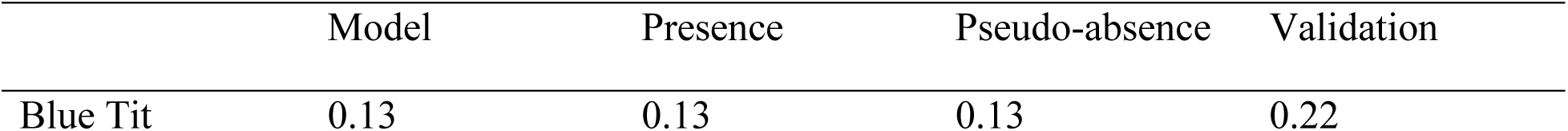
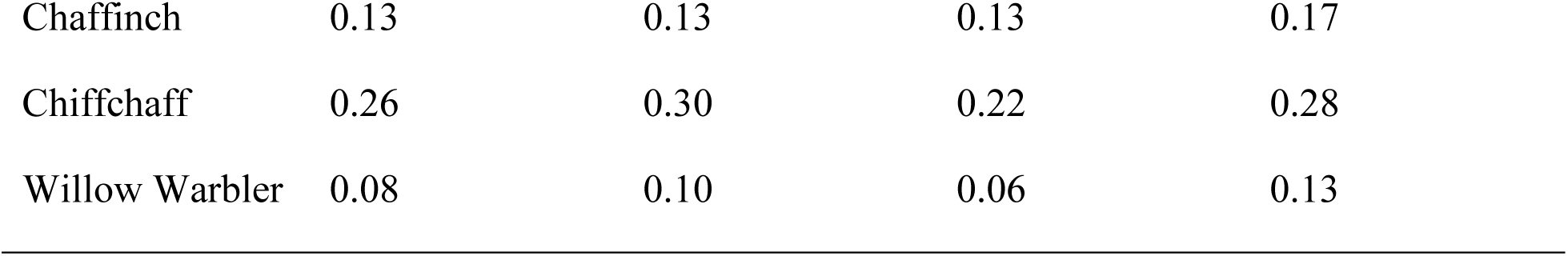
Model, presence and pseudo-absence out-of-bag (OOB) error and validation error for RF models developed with Landsat data.

|  | Model | Presence | Pseudo-absence | Validation |
| --- | --- | --- | --- | --- |
| Blue Tit | 0.13 | 0.13 | 0.13 | 0.22 |
| Chaffinch | 0.13 | 0.13 | 0.13 | 0.17 |
| Chiffchaff | 0.26 | 0.30 | 0.22 | 0.28 |
| Willow Warbler | 0.08 | 0.10 | 0.06 | 0.13 |

Results from the ALS RF models are summarized in Table 11. The lowest model error was obtained for Willow Warbler, followed by Chaffinch, Blue Tit and Chiffchaff. Validation error had the same pattern and error values were equal or slightly greater. Model, presence and pseudo-absence OOB errors were equal for Willow Warbler and similar for Blue Tit. Chaffinch and Chiffchaff both had lower pseudo-absence error. Relative to the corresponding Landsat models for all species, the OOB model and pseudo-absence error is greater with ALS RF models, and OOB presence error is equal or greater. Validation error displays a similar pattern with slightly higher error for all species except for Willow Warbler which has slightly lower error in the ALS models.

**Table 11.** Model, presence and pseudo-absence out-of-bag (OOB) error and validation error for RF models developed with ALS data.

|  | Model | Presence | Pseudo-absence | Validation |
| --- | --- | --- | --- | --- |
| Blue Tit | 0.22 | 0.22 | 0.21 | 0.23 |
| Chaffinch | 0.19 | 0.25 | 0.13 | 0.19 |
| Chiffchaff | 0.29 | 0.35 | 0.24 | 0.31 |
| Willow Warbler | 0.10 | 0.10 | 0.10 | 0.10 |

The error for RF models developed with multi-sensor fusion combining ALS and Landsat data are described in Table 12. The lowest OOB model error was obtained for Willow Warbler, followed by Chaffinch, Blue Tit and Chiffchaff. OOB model, presence and pseudo-absence error values were equal or similar for Willow Warbler, Chaffinch and Blue Tit. OOB presence error was greater than pseudo-absence error for Chiffchaff. Validation error was smallest for Willow Warbler, followed by Blue Tit, Chaffinch and Chiffchaff. The fusion OOB model error values for all species are lower than those from RF ALS models, and lower than RF Landsat models for Chiffchaff and Willow Warbler. Validation errors are lower than both ALS and Landsat RF models for all bird species.

**Table 12.** Model, presence and pseudo-absence out-of-bag (OOB) error and validation error for RF models developed with multi-sensor fusion combining ALS and Landsat data.

|  | Model | Presence | Pseudo-absence | Validation |
| --- | --- | --- | --- | --- |
| Blue Tit | 0.15 | 0.14 | 0.15 | 0.09 |
| Chaffinch | 0.14 | 0.14 | 0.14 | 0.14 |
| Chiffchaff | 0.20 | 0.23 | 0.17 | 0.25 |
| Willow Warbler | 0.07 | 0.07 | 0.07 | 0.06 |

Finally, the error for RF models developed with the predicted surfaces from our first objective are described in Table 13. For all species, pseudo-absence error was lower than presence error. OOB model and validation error values were lowest for Willow Warbler, followed by Chaffinch, Blue Tit and Chiffchaff, a similar pattern as previous models. However, the values for OOB model, presence, pseudo-absence error and validation error were higher relative to all other models for all species.

**Table 13.** Model, presence and pseudo-absence out-of-bag (OOB) error and validation error for RF models developed with predicted surfaces generated through multi-sensor fusion of structural attributes with spectral data.

|  | Model | Presence | Pseudo-absence | Validation |
| --- | --- | --- | --- | --- |
| Blue Tit | 0.24 | 0.27 | 0.21 | 0.33 |
| Chaffinch | 0.21 | 0.26 | 0.17 | 0.28 |
| Chiffchaff | 0.32 | 0.38 | 0.27 | 0.47 |
| Willow Warbler | 0.18 | 0.19 | 0.17 | 0.24 |

The variable importance plots for Landsat, ALS and data fusion RF models (Figures 2-5) show the MDA for each variable. MDA values represent the decline in accuracy that would occur if that variable was removed from the models. Variables are plotted in descending order of importance for ease of interpretation.

**Figure 2.**
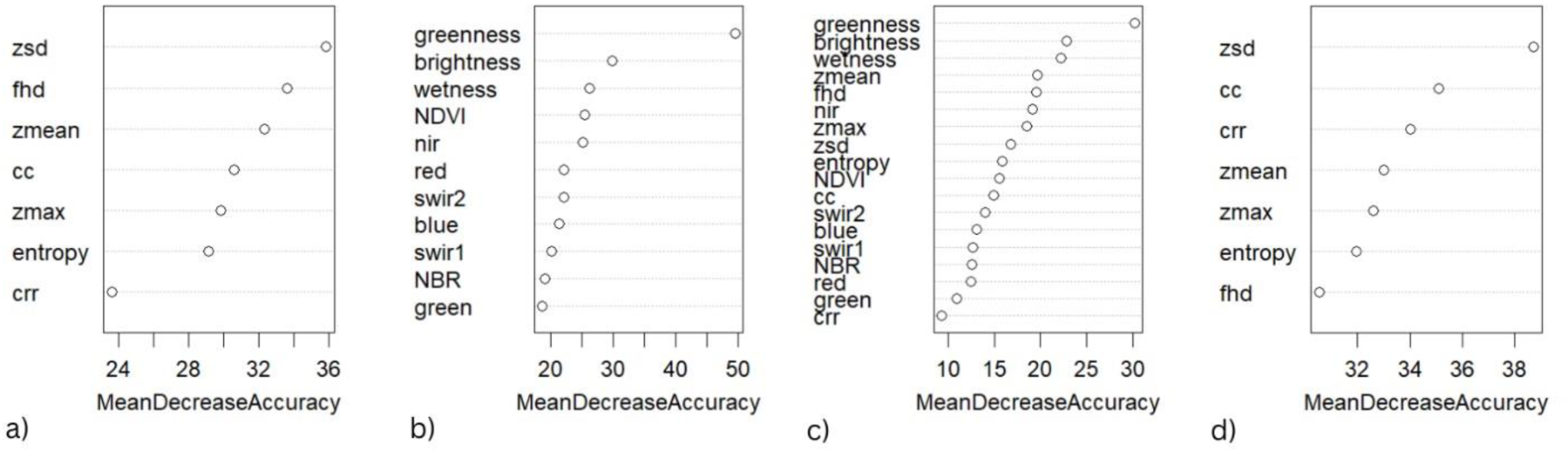
Variable importance for Blue Tit from RF models developed with a) ALS data, b) Landsat data c) object-level fusion data from objective 2, and d) predicted structural surfaces from objective 1.

For Blue Tit (Figure 2), the most important variables in the Landsat model and in the multisensor fusion model combining ALS and Landsat data were TCT brightness, greenness and wetness. For the ALS model, the top variables were standard deviation of height, FHD and mean height.

Standard deviation of heights was also the most important in the model developed with predicted surfaces, followed by CC and CRR. The greatest decrease in accuracy was for TCT greenness in the Landsat model (and object-level fusion model).

For Chaffinch (Figure 3), maximum, mean and standard deviation of heights were the most important in the ALS model. TCT brightness, greenness and wetness were most important in the Landsat model. In the object-level fusion model for Chaffinch, Landsat TCT greenness and wetness and ALS maximum height were the most important variables. In the prediction surface model, standard deviation of heights, maximum height and canopy cover were the most important variables. Across all models, the greatest loss in accuracy with removal was for standard deviation of heights in the model developed with predicted surfaces.

**Figure 3.**
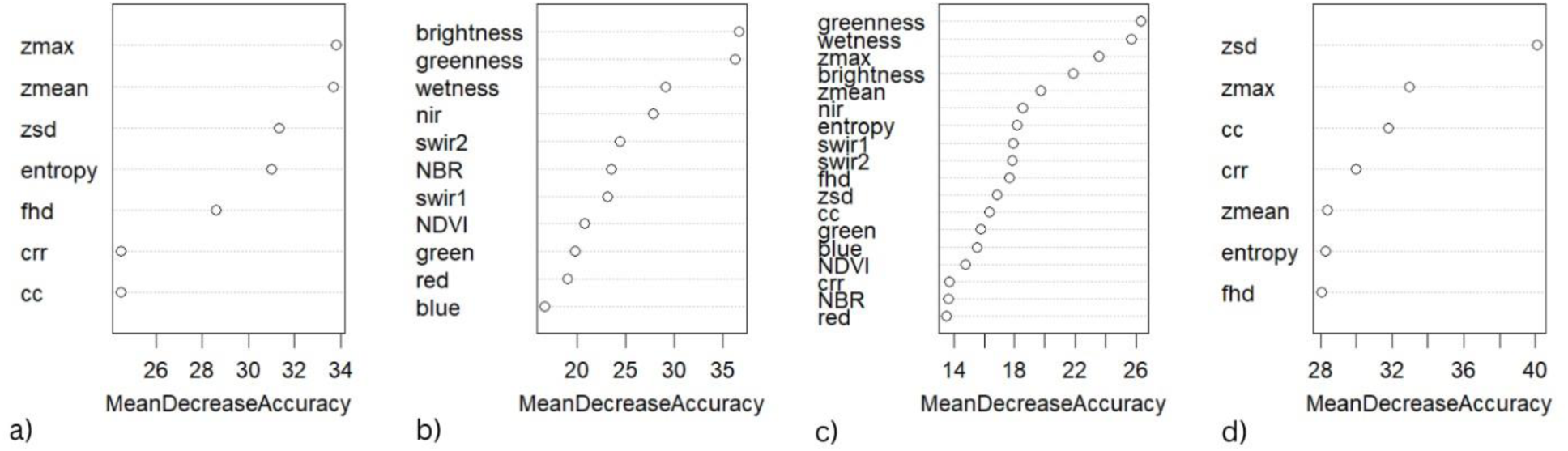
Variable importance for Chaffinch from RF models developed with a) ALS data, b) Landsat data c) object-level fusion data from objective 2, and d) predicted structural surfaces from objective 1.

For Chiffchaff (Figure 4), the most important variables in the ALS model were maximum, mean, and standard deviation of heights. In the Landsat model, TCT brightness and wetness, and information from the SWIR 2 band were most important. In the object-level fusion model, the top variables were maximum and mean height from ALS and TCT brightness from Landsat. The most important variables in the prediction surface model were standard deviation of heights, FHD and CC. The greatest loss of accuracy with removal was for maximum height in the ALS model.

**Figure 4.**
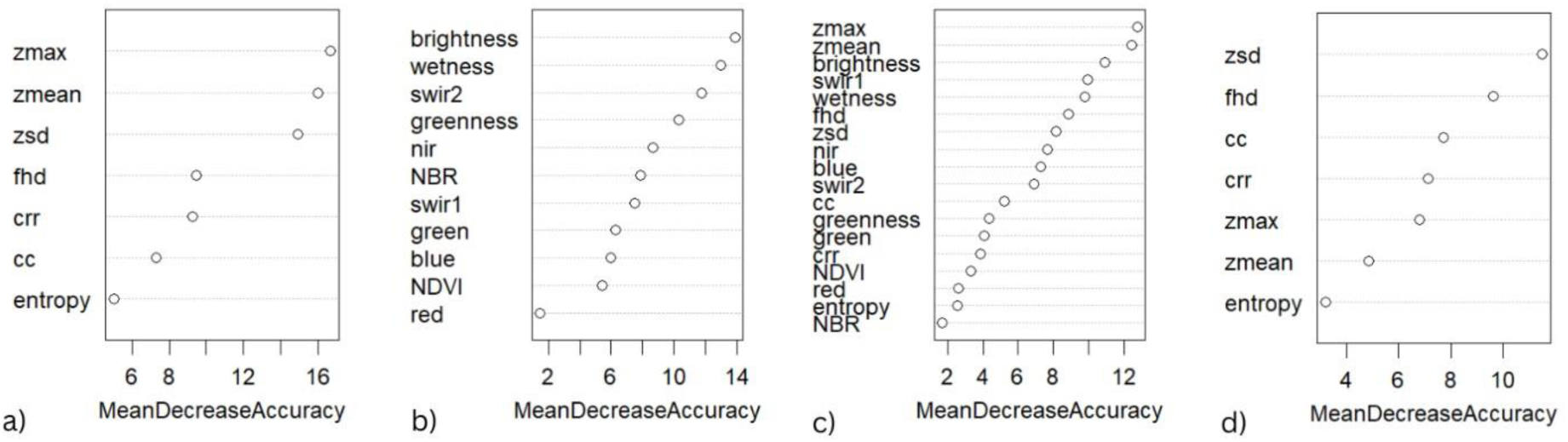
Variable importance for Chiffchaff from RF models developed with a) ALS data, b) Landsat data c) object-level fusion data from objective 2, and d) predicted structural surfaces from objective 1.

Finally, for Willow Warbler (Figure 5), the most important variables in the ALS model were standard deviation of heights, maximum and mean height. TCT wetness, brightness and NDVI were the most important in the Landsat model. In the object-level fusion model, Landsat-derived TCT wetness and ALS-derived standard deviation of heights and maximum height were the most important. Maximum height, standard deviation of heights and entropy were identified as the most important in the prediction surface model. Removal of the standard deviation of heights in the ALS model would result in the greatest loss in accuracy.

**Figure 5.**
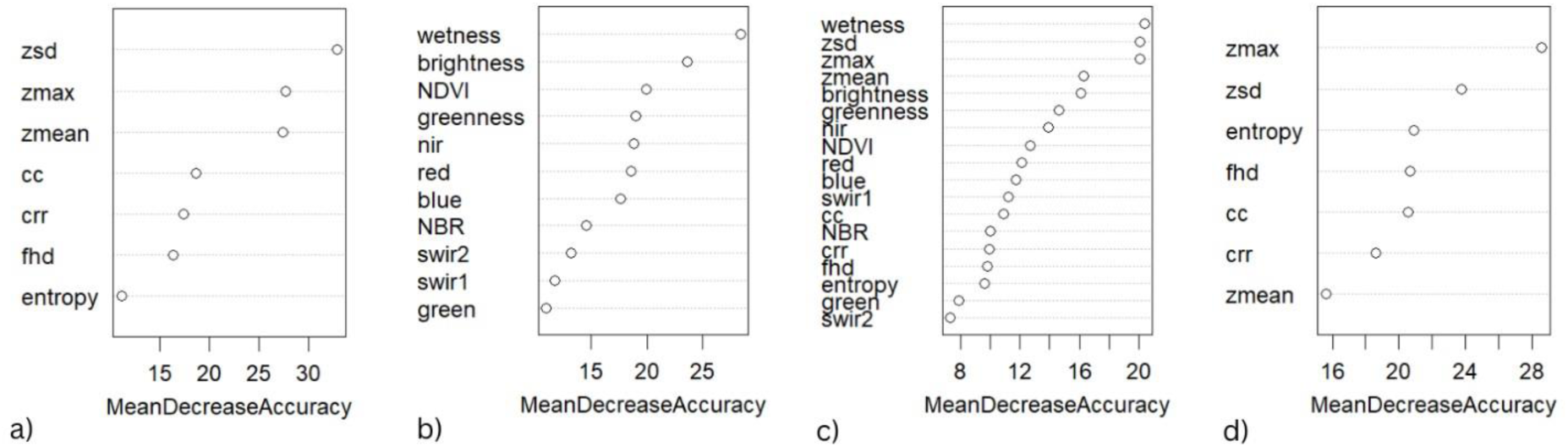
Variable importance for Willow Warbler from RF models developed with a) ALS data, b) Landsat data c) object-level fusion data from objective 2, and d) predicted structural surfaces from objective 1.

Some variables were identified as important for multiple species. In the ALS model, standard deviation of heights and mean height were among the top three most important variables for Blue Tit, Chaffinch, Chiffchaff and Willow Warbler. Maximum height was important for all but Blue Tit. In the Landsat models, TCT brightness and wetness were important to all species. TCT greenness was important to Blue Tit and Chaffinch. In the object-level fusion models, TCT brightness (Blue Tit, Chiffchaff), TCT greenness (Blue Tit, Chaffinch), TCT wetness (Blue Tit, Chaffinch, Willow Warbler), and ALS-derived maximum height (Chaffinch, Chiffchaff, Willow Warbler), were repeated across multiple species though no variables were shared among the top three for all four bird species. The shapes of the relationships between species occurrence and important variables shared across species are given in supplementary material (Figures S8 and S9).

### 3.3 Predicting bird habitat

Prediction error based on spatial intersection of actual presence sampling plot locations with predicted presence pixels is summarized in Table 14. For Blue Tit and Willow Warbler, presence habitat prediction error was low for the ALS model on ALS data, Landsat model on Landsat data and object-level fusion model on fused data. For Chaffinch and Chiffchaff, the lowest error was for the ALS model on ALS data, followed by the fusion model on the fused data, and the Landsat model on the Landsat data. Except for Chaffinch, the highest error was associated with either the ALS model on the prediction surface or the prediction surface model on the prediction surface.

**Table 14.** Prediction error associated with Blue Tit, Chaffinch, Chiffchaff and Willow Warbler presence locations at Old Wilderness and New Wilderness for the five prediction (P) scenarios (described in Table 5).

|  | P_ALS | P_Landsat | P_Comb | P_Pred | P_ALSandPred |
| --- | --- | --- | --- | --- | --- |
| Blue Tit | 0.01 | 0.02 | 0.02 | 0.33 | 0.38 |
| Chaffinch | 0.09 | 0.20 | 0.16 | 0.24 | 0.15 |
| Chiffchaff | 0.14 | 0.38 | 0.26 | 0.56 | 0.38 |
| Willow Warbler | 0.03 | 0.01 | 0.01 | 0.09 | 0.15 |

We produced a series of surfaces (Figure 6) for the five prediction scenarios for each of the four study species at the Old Wilderness study site in 2015. Pixels are classified as presence (yellow) or pseudo-absence (purple), and the actual presence sampling plots are overlaid (white circles).

**Figure 6.**
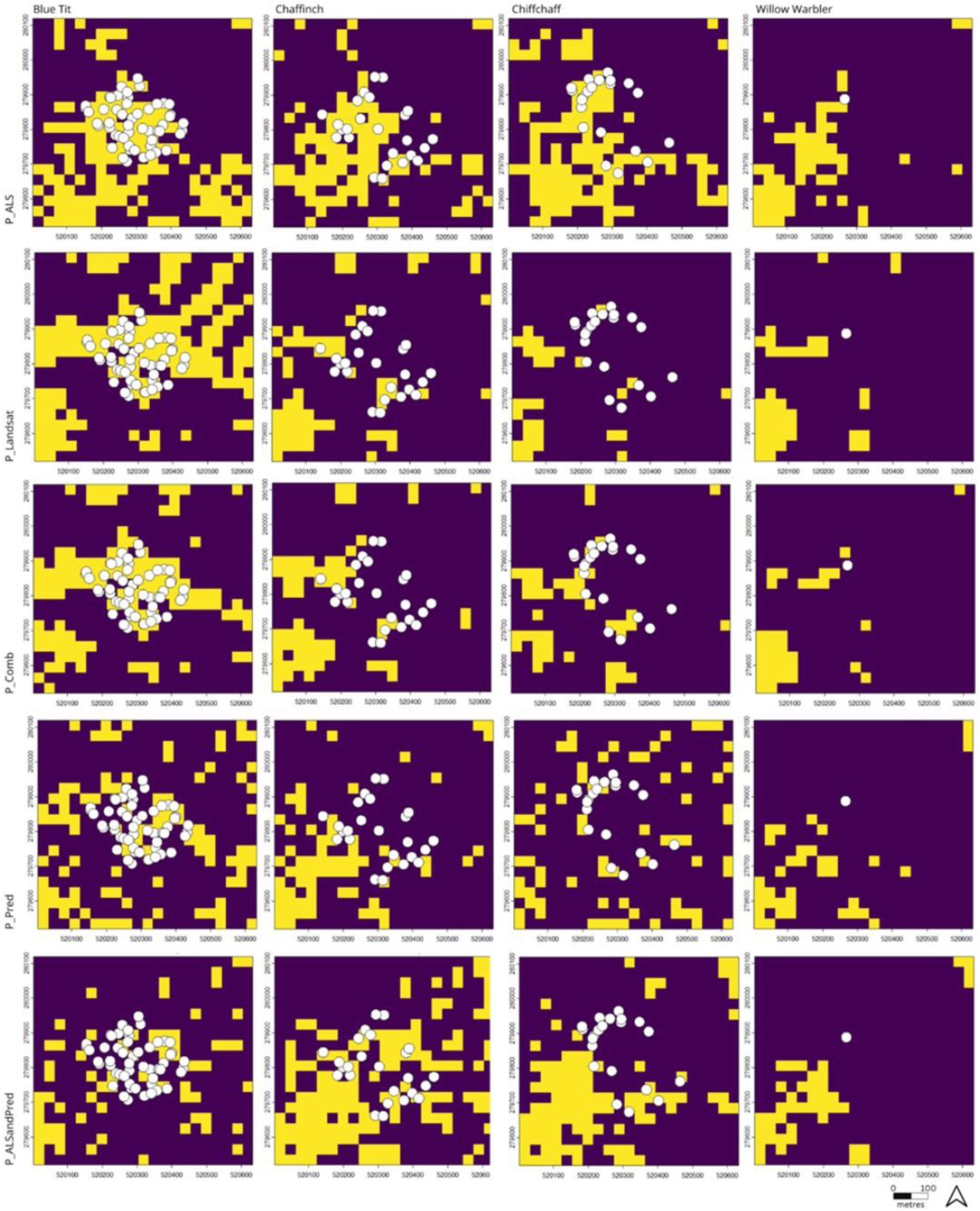
Prediction surfaces for Blue Tit (first column), Chaffinch (second column), Chiffchaff (third column), and Willow Warbler (fourth column) at the Old Wilderness for the five prediction scenarios described in Table 5: P_ ALS (first row), P_Landsat (second row), P_ Comb (third row), P_ Pred (fourth row) and P_ALSandPred (fifth row). Yellow pixels represent presence, purple pixels represent pseudo-absence. Pixels are 30 x 30 metres.

Blue Tit and Willow Warbler had low prediction error for ALS only, Landsat only, and objectlevel fusion scenarios and, as shown in Figure 6, within the Old Wilderness study site these three surfaces largely agree. Blue Tit occurs across the Old Wilderness, and pixels within this study site are largely predicted as areas of presence. Willow Warbler, on the other hand, has limited presence in the Old Wilderness later in the time series with only one presence recorded, and much of the study site was predicted as areas of absence. Both prediction scenarios that use the predicted structural attributes from our first objective underestimate presence, and presence pixels are more fragmented for Blue Tit. In all prediction scenarios for Willow Warbler, presence areas were limited, while scenarios that use the predicted surfaces from our first objective also predict less presence pixels, although the magnitude of difference is not as great because there was less presence predicted overall.

Chaffinch and Chiffchaff had the lowest error for the ALS models. For both species, most predictions underestimated presence pixels, except for the prediction scenario using only predicted structural surfaces. Many of the actual presences for Chaffinch and Chiffchaff occur along the perimeter of the Old Wilderness, with fewer presences recorded in the centre of this study site. Here again, predictions using solely the predicted surfaces from the first objective had low accuracy. Interestingly, the ALS model predicting on this surface performed reasonably well relative to other prediction scenarios for each species, particularly for Chaffinch. Compared to Blue Tit and Willow Warbler, the spatial predictions between this scenario and the ALS predictions largely agree and identify similar presence pixels.

Overall, the more spatially widespread species (Blue Tit) and spatially restricted species (Willow Warbler) within our study sites were the most accurately predicted. Furthermore, predictions with ALS, Landsat, and object-level fusion generally preformed equally well and agreed on presence pixel predictions within our study site. For Chaffinch and Chiffchaff, which tended to be present along the edges of our study sites, the ALS predictions were more accurate despite higher model performance for Landsat models relative to ALS models. The Landsat models underpredicted presence pixels relative to the ALS predictions.

## 4 Discussion

This study used multi-temporal, multi-sensor data to systematically implement two fusion methods with spectral and structural remote sensing variables derived from Landsat and ALS data. First, we implemented a pixel-level approach to predict multiple structural attributes using Landsat spectral data. Second, we adopted an object-level approach to develop SDMs for Blue Tit, Chaffinch, Chiffchaff and Willow Warbler with the ALS data, Landsat data, object-level data fusion, and the predicted structural attributes. Third, we predicted future presence for each bird species using the developed SDMs and the ALS data, Landsat data, object-level data fusion, and structural attributes predicted with pixel-level fusion. The objective of both fusion methods was to assess the ability of Landsat and ALS data and fusion for modelling and predicting habitat.

Previous studies have used the relationship between spectral and structural remote sensing data to predict individual attributes (e.g., canopy height; Staben et al., 2018; Wilkes et al., 2015) or multiple measured and derived forest structural attributes simultaneously (Matasci et al., 2018a), across broader spatiotemporal extents. Upscaling may be useful in applications ranging from carbon modelling (e.g., upscaling ALS-derived above ground biomass estimates with Landsat; Hudak et al., 2020) to habitat modelling (e.g., upscaling GEDI with Landsat; Smith et al., 2022). We were able to model multiple structural attributes simultaneously through pixel-level fusion with Landsat data and found that information from the green band and NBR were consistently ranked as important, except for entropy which had high OOB error. Contrary to other studies (e.g., Ahmed et al., 2015), when we examined how error was manifested across space it was higher in mature rather than young forest environments. Validation data were from the same years that were used to train the models, whereas the prediction data were entirely omitted from model development. As expected, RMSE values on the prediction data were slightly higher than the validation dataset. While the magnitude of these prediction errors (e.g., differences ranging from 2.55 to 3.86 metres for mean height) may be acceptable for some applications, and even for modelling and predicting the habitat of some bird species, these errors were likely to impact SDM accuracy. We anticipated that this error would be magnified when used in SDM development, and when these predicted structural attributes are used in SDM for specialist bird species that use a narrow range of habitat structure.

Indeed, for all bird species, SDMs developed with the predicted surfaces had the highest OOB model and validation error. Distinct structural conditions that are important to bird habitat may have similar spectral attributes (Adams & Matthews, 2018), contributing to this confusion in prediction. Based on OOB and validation error values, the Landsat SDMs were best for Blue Tit and Chaffinch, while the object-level fusion models were best for Chiffchaff and Willow Warbler. Species-specific traits such as habitat associations and generality are important to modelling outcomes. The lower accuracy of ALS models for the generalist species, Blue Tit and Chaffinch, is likely due to the diverse vegetation structures that they occupy and the spatial aggregation of ALS data to 30 metre resolution which contributed to the compressed range of the output variables. Landsat derived spectral characteristics sufficiently characterized the habitat of these two generalists. Habitat specialists, like Chiffchaff and Willow Warbler, occupy a narrower range of environmental conditions (Devictor et al., 2010). For instance, Willow Warbler favours early successional forest (Saether, 1983) which has a distinct structure that ALS data can be used to characterize. However, the ALS models for Chiffchaff and Willow Warbler were less accurate than the Landsat models, but ALS data created more accurate models when fused with Landsat data. Our results suggest that data choice (e.g. spectral, structural, spatial resolution, etc.) should be aligned with the ecology of the study species. Previous studies have noted higher accuracies through remote sensing data fusion (e.g., with radar, lidar, and spectral data in Swatantran et al., 2012), which is aligned with the accuracy gains for Willow Warbler and Chiffchaff. Contrary to our expectation, object-level fusion did not outperform Landsat and ALS models for all species, and for Blue Tit and Chaffinch data fusion only added confusion.

The MDA values provided by RF identified many shared variables that were important to multiple species. Maximum, mean and standard deviation heights from ALS, and TCT brightness, greenness and wetness were highly ranked across multiple species across the ALS, Landsat and object-level fusion models. These ALS variables have been identified in many other bird habitat studies (see review by Bakx et al., 2019). For instance, maximum and mean heights derived from ALS data were important not just to presence, but to nestling body mass (Hill & Hinsley, 2015). The high importance of FHD for Blue Tit only was surprising given that this attribute has a long history in bird habitat studies (e.g., since MacArthur & MacArthur, 1961) and has been found to be important in other studies, particularly studies of species diversity rather than single species occurrence (i.e., FHD was important to bird diversity in Melin et al., 2019). It was surprising that TCT variables were highly ranked given that these are not extensively used in previous bird habitat studies (with some exceptions, e.g., TCT variables have been found to be significant in Alessandrini et al., 2022; Moreira et al., 2022) but are more widely used in the remote sensing literature.

From a modelling standpoint, in years where both ALS and Landsat data are available, an object-level fusion approach to model habitat is recommended to more accurately characterize the habitat of specialists like Chiffchaff and Willow Warbler. If only ALS or Landsat data are available, either remote sensing data type can be used in SDM development to produce accurate results for generalist species, like Blue Tit and Chaffinch. In years when no ALS data are available, our results suggest that the spectral information and derived metrics from Landsat data may be more suitable than using Landsat to predict structural attributes. Regardless of the data used or fused, candidate predictor variables should be identified using ecological knowledge for the study species and refined through the model fitting process.

While RF MDA can be used to rank or identify important variables, it does not provide the values associated with presence for the study species (but see distribution of values at presence locations for all species in the supplementary materials; Figures S8 and S9). Within the New Wilderness and Old Wilderness, Blue Tit presence was associated with higher maximum heights, a range of mean height values and intermediate standard deviation of heights. Chaffinch presence was characterized by higher maximum height values than Blue Tit, intermediate mean height values, and similar standard deviation of heights values, with a smaller second peak at higher values. Chiffchaff occurs across a range of maximum height values, but lower mean and standard deviation of height values than either Blue Tit or Chaffinch. Willow Warbler was notably associated with lower values for all three ALS variables. For the TCT variables, all species show a similar bimodal pattern for brightness but Willow Warbler has a larger peak at greater values. For TCT greenness, Blue Tit and Chiffchaff have a peak at higher values, while Chaffinch and Willow Warbler have a more bimodal distribution with a higher peak at greater TCT greenness for Chaffinch and a higher peak at lower values for Willow Warbler. Finally, for TCT wetness, Willow Warbler presence is associated with higher values, Blue Tit and Chiffchaff presences are characterized by lower values, and Chaffinch presence was associated with a wider range of intermediate values. The Landsat and ALS variables capture ecologically meaningful information about the study species’ habitat. For instance, as noted above, Willow Warbler favours early successional environments which can be characterized by the occurrence of shrubby vegetation that is shorter and less variable in stature (e.g., lower maximum and standard deviation of height values) and more open areas (e.g., higher TCT brightness and lower TCT greenness values). Note that there were differences across lower ranked variables, here we focused on the shared important variables to compare across species (Figures S8 and S9).

ALS and spectral data fusion for the prediction of forest structural attributes has become established, particularly in forest management applications for the estimation of inventory attributes such as basal area, volume and biomass (Lim et al., 2003; Matasci et al., 2018a; Matasci et al., 2018b; Wulder et al., 2012). While fusion for prediction of structural attributes is less explored in wildlife and biodiversity applications, continuous spatiotemporal prediction of structural attributes is important (Vogeler et al., 2023). Furthermore, from a management, monitoring or conservation perspective, it may be useful to predict structural habitat attributes due to their ease of interpretation. For instance, maximum height from ALS is likely to be more readily understood by conservation officers and operationalized in conservation policies than Landsat-derived TCT variables. However, our results suggest that pixel-level fusion to predict ALS structural attributes with Landsat spectral data may not sufficiently capture relevant habitat characteristics. As expected, we found that presence prediction error was generally greatest when predicted surfaces were used. Further research should consider the impacts of model selection, variable selection and extraction, study species, and the scale and resolution of analysis to improve the spatiotemporal prediction of structural attributes. For instance, Landsat variables were extracted using an area-weighted mean and this may have contributed to the low prediction accuracy for Chaffinch and Chiffchaff with Landsat data. These species were present along the edges of the study sites, and a mean value from multiple pixels across a forest edge will not adequately capture the relevant information of that important feature. For modelling, we found a distinction between generalist and specialist species. While for prediction, there is a distinction between edge and interior species. Interestingly, the interior species also represent the most spatially widespread and abundant (Blue Tit) and restricted and uncommon (Willow Warbler) species. However, this may reflect the particular distribution of these species preferred habitat at our study sites.

Through our two fusion approaches, we were able to predict structural attributes with spectral data and improve the characterization of habitat for Chiffchaff and Willow Warbler. However, the error associated with structural attribute prediction was magnified when these surfaces were used to model and predict habitat, and not all species benefited from object-level fusion. Models may be improved further by considering other Landsat and ALS variables, by including higher spatial resolution remote sensing data, and by comparing other modelling methods to the performance of RF.

First, accuracy may also be improved through the inclusion of other variables. There is ongoing debate in the habitat modelling community regarding variable selection in models. One view promotes limiting the number of variables based on existing ecological knowledge and study goals (Elith & Leathwick, 2009), while an alternate view is to retain all information (e.g., “fit first, explain later” in Ciuti et al., 2018). There are merits to both philosophies, and there is an expansive number of variables that can be extracted from ALS point clouds and Landsat spectral data. We limited the variables in this study to include those that are commonly used (e.g., NDVI), that we expected may be important (i.e., FHD), and that describe complementary aspects of habitat (i.e., TCT brightness, greenness and wetness, horizontal and vertical heterogeneity). It is possible that other variables are important for characterizing the breeding season habitat of Blue Tit, Chaffinch, Chiffchaff and Willow Warbler, and for predicting ALS structural forest characteristics from Landsat spectral data. For instance, we only used Landsat data that were available from both the TM and OLI sensors (i.e., blue, green, red, NIR, SWIR 1, SWIR 2), so the higher resolution OLI panchromatic band was not used directly or for pansharpening (Anand & Sharma, 2024). Finer spatial information through pansharpening may contribute to increased prediction of the structural variables, which may in turn reduce the error propagation when predicted surfaces are used in SDMs. Pansharpening has resulted in higher accuracies of bird population density (Richter et al., 2020) and tree species classification (Deur et al., 2021). However, contrasting results have also been found where coarser resolution data yielded higher accuracy in SDMs (Wunderlich et al., 2022), thereby warranting further examination. Furthermore, variables describing relevant vegetation layers (e.g., overstorey, understorey, shrub) can be extracted from 3D ALS point clouds. ALS data were not stratified in the present study to maintain consistency in the variables used for both fusion approaches as we would not expect Landsat spectral data to be able to predict these types of variables. However, the lower layers in the canopy are associated with greater avian diversity (e.g., vegetation below 4 metres in Melin et al., 2018) and variables describing vegetation layers have been found to be important (e.g., for Blue Tit in *Kuzmich forthcoming*).

Next, as suggested above, spectral remote sensing data with a higher spatial resolution would likely improve accuracy. We chose to use Landsat data because it is openly available, it temporally covered the full ALS time series (albeit with different Landsat sensors) and is commonly used in habitat studies. It would be worthwhile to test any gains in accuracy with higher resolution imagery. For instance, an unmanned aerial vehicle (UAV) could be deployed routinely at both study sites to capture data at a significantly higher spatial resolution (e.g., Dandois & Ellis, 2013). Similarly, high spatial resolution satellite data such as Sentinel-2 may improve accuracy as measures captured using a smaller grain size have been found to be more strongly correlated with ALS than the same measures with a larger grain size (e.g., texture metrics with a 10 metre resolution versus 30 metre resolution; Farwell et al., 2021).

Finally, other modelling forms may have yielded more accurate results as machine learning methods such as RF have been reported to have lower transferability in some scenarios (i.e., Jin et al., 2018). While our modelling scenarios considered temporally proximate years, it is unclear to what extent they would be generalizable through time. Our SDMs address this issue by including variables from both study sites, which represent different successional stages, across multiple years so that the data used for training/validation and temporal cross-validation were similar. We also chose RF for the object-level data fusion to model habitat and the pixel-level data fusion to predict ALS structural forest characteristics so that the modelling method did not bias our results. RF methods are able to fit complex relationships, which may limit their predictive ability at sites that are dissimilar from the sites used to develop the models. It would be worthwhile for future studies to examine differences across modelling methods.

## 5 Conclusion

This study systematically assessed two fusion methods; a pixel-based method to predict structural attributes from spectral data and an object-level method to model bird habitat. The implicit question was to determine how best to use or fuse ALS structural and Landsat spectral data and derived attributes to model bird habitat. Through our pixel-based fusion, we found that ALS structural attributes could be predicted from spectral data but that the error was propagated and intensified in SDMs. Through our object-level fusion to develop SDMs, we found that habitat specialists like Chiffchaff and Willow Warbler had higher accuracy than ALS or Landsat alone, but that this was not the case for generalists like Blue Tit and Chaffinch. This study contributes to remote sensing data fusion and habitat modelling by systematically testing how multi-temporal Landsat and ALS data can be used to model habitat for two dynamic young forest study sites. Future consideration should be given to improve our understanding of the relationship between spectral and structural data in the context of habitat modelling with care given to the ecology of the study species, which we anticipate will translate to improved accuracy.

## Acknowledgements

Data collection was supported by the UK Centre for Ecology and Hydrology, the Royal Society for the Protection of Birds, the UK Environment Agency, the Natural Environment Research Council Airborne Research and Survey Facility, Bournemouth University, and Queen’s University. We also wish to acknowledge R. Broughton and P. Davies for their contributions to bird data collection and digitization.

## CRediT co-authorship

Rachel J Kuzmich: Writing – original draft, Funding acquisition, Conceptualization, Methodology, Project administration, Formal Analysis, Investigation, Data curation, Software, Visualization, Validation. Ross A Hill: Writing – review & editing, Supervision, Methodology, Validation, Data curation, Resources. Shelley A Hinsley: Writing – review & editing, Investigation, Methodology, Project administration, Data curation, Resources. Paul E Bellamy: Writing – review & editing, Investigation, Methodology, Project administration, Data curation, Resources. Ailidh E Barnes: Writing – review & editing, Investigation, Data curation, Resources. Markus Melin: Writing – review & editing, Investigation, Data curation, Resources. Paul M Treitz: Writing – review & editing, Supervision, Funding acquisition, Validation, Resources.

## Data availability

Remote sensing datasets are available via the United States Geological Survey (Landsat) and the Centre for Environmental Data Analysis (ALS). Bird data are available upon reasonable request to Rachel J Kuzmich. R code is available at https://github.com/rachelkuzmich/multi_temp_and_sensor_bird_habitat

## Supplementary Materials

**S1 Figure:**
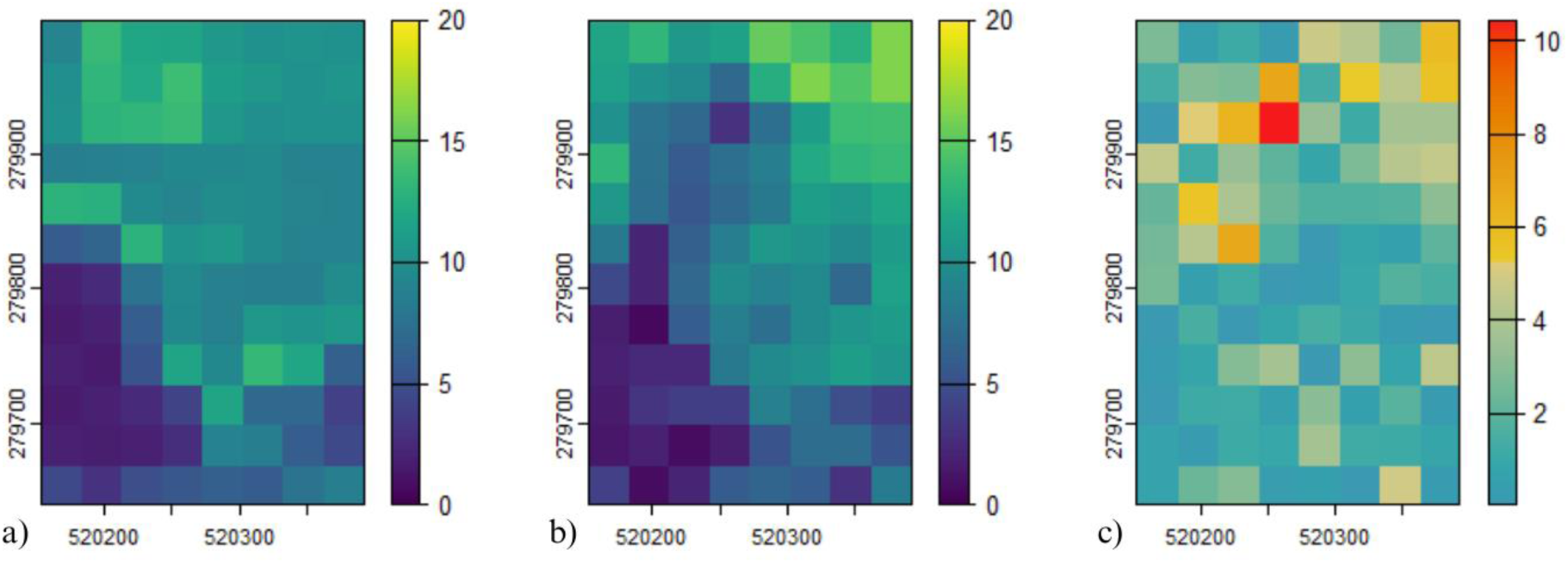
predicted, actual and differenced surfaces for maximum height, S2 Figure: predicted, actual and differenced surfaces for mean height, S3 Figure: predicted, actual and differenced surfaces for standard deviation of heights, S4 Figure: predicted, actual and differenced surfaces for entropy, S5 Figure: predicted, actual and differenced surfaces for foliage height diversity, S6 Figure: predicted, actual and differenced surfaces for canopy cover, S7 Figure: predicted, actual and differenced surfaces for canopy relief ratio, S8 Figure: habitat characteristics for important ALS variables, and S9 Figure: habitat characteristics for important Landsat variables. S1 Figure. Surfaces depicting the a) predicted maximum height values from Random Forests for 2015, b) rasterized ALS data for 2015, and c) a difference raster. All values are in metres. Pixels are 30 x 30 metres.

**S2 Figure.**
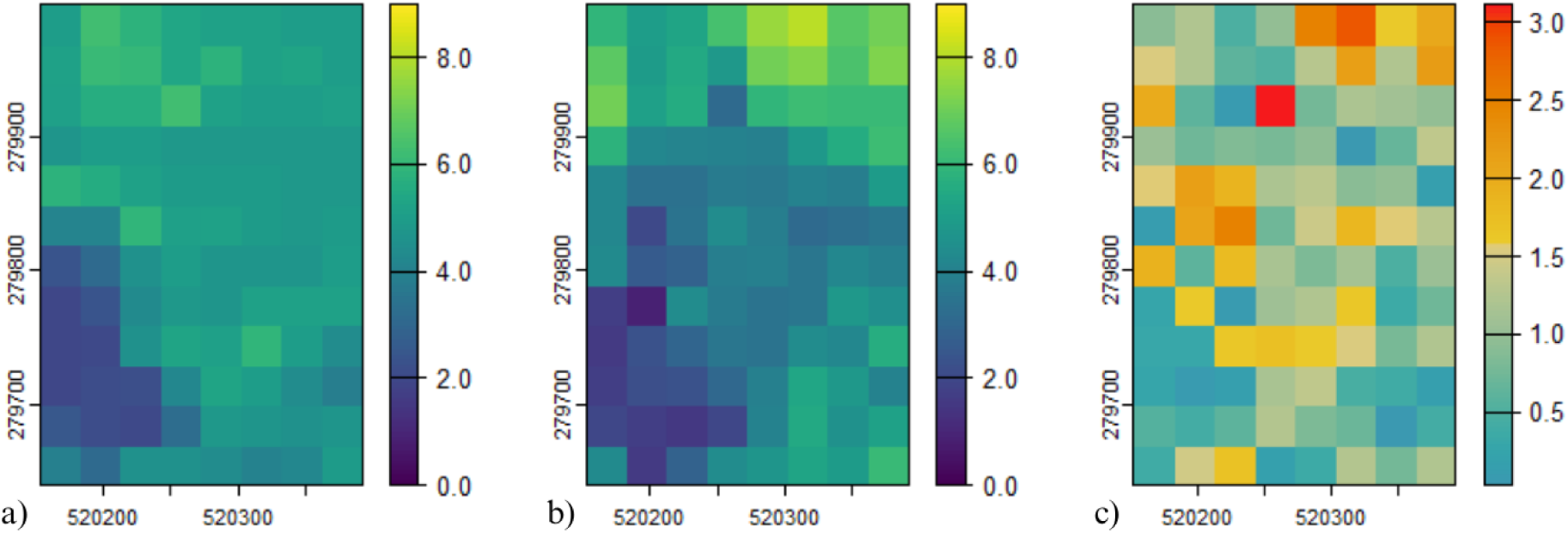
Surfaces depicting the a) predicted mean height values from Random Forests for 2015, b) rasterized ALS data for 2015, and c) a difference raster. All values are in metres. Pixels are 30 x 30 metres.

**S3 Figure.**
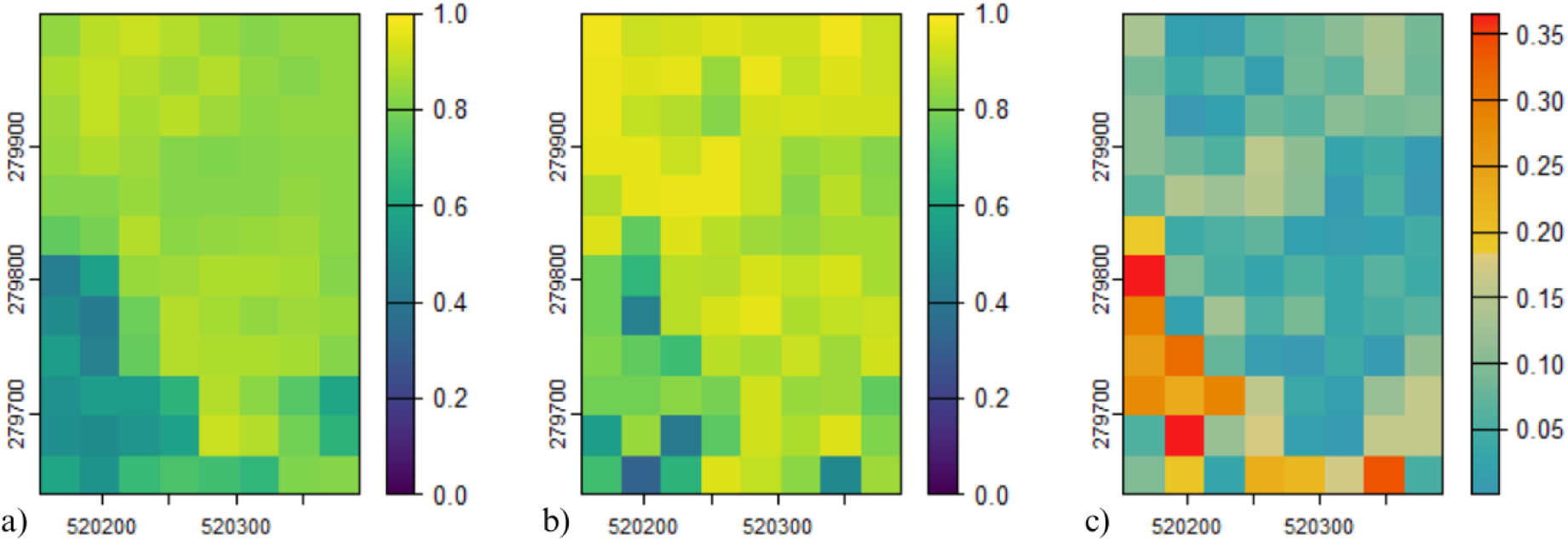
Surfaces depicting the a) predicted standard deviation of height values from Random Forests for 2015, b) rasterized ALS data for 2015, and c) a difference raster. All values are in metres. Pixels are 30 x 30 metres.

**S4 Figure.**
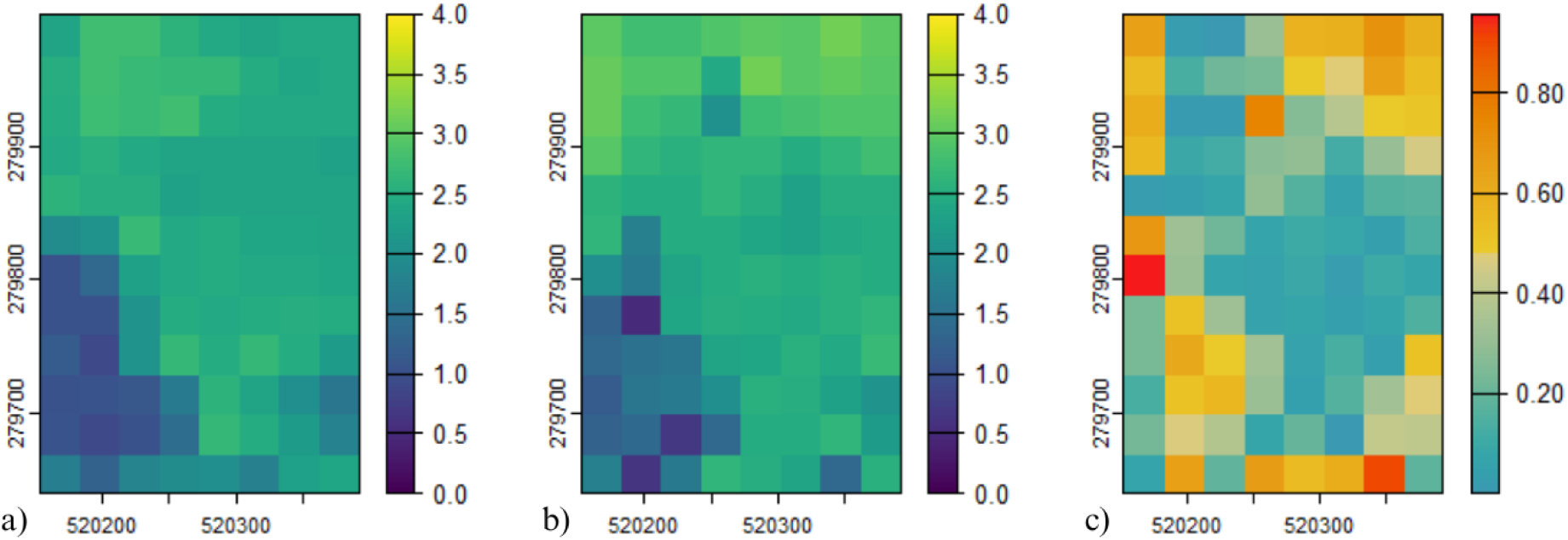
Surfaces depicting the a) predicted entropy values from Random Forests for 2015, b) rasterized ALS data for 2015, and c) a difference raster. Pixels are 30 x 30 metres.

**S5 Figure.**
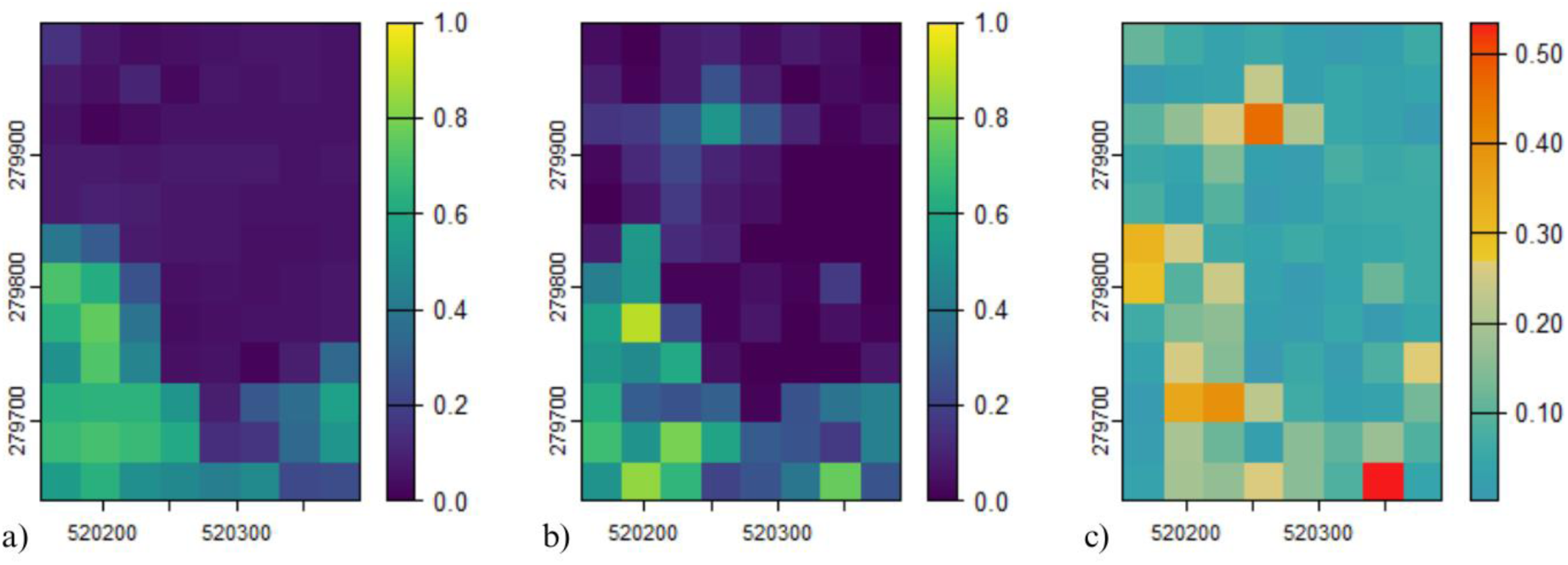
Surfaces depicting the a) predicted FHD values from Random Forests for 2015, b) rasterized ALS data for 2015, and c) a difference raster. Pixels are 30 x 30 metres.

**S6 Figure.**
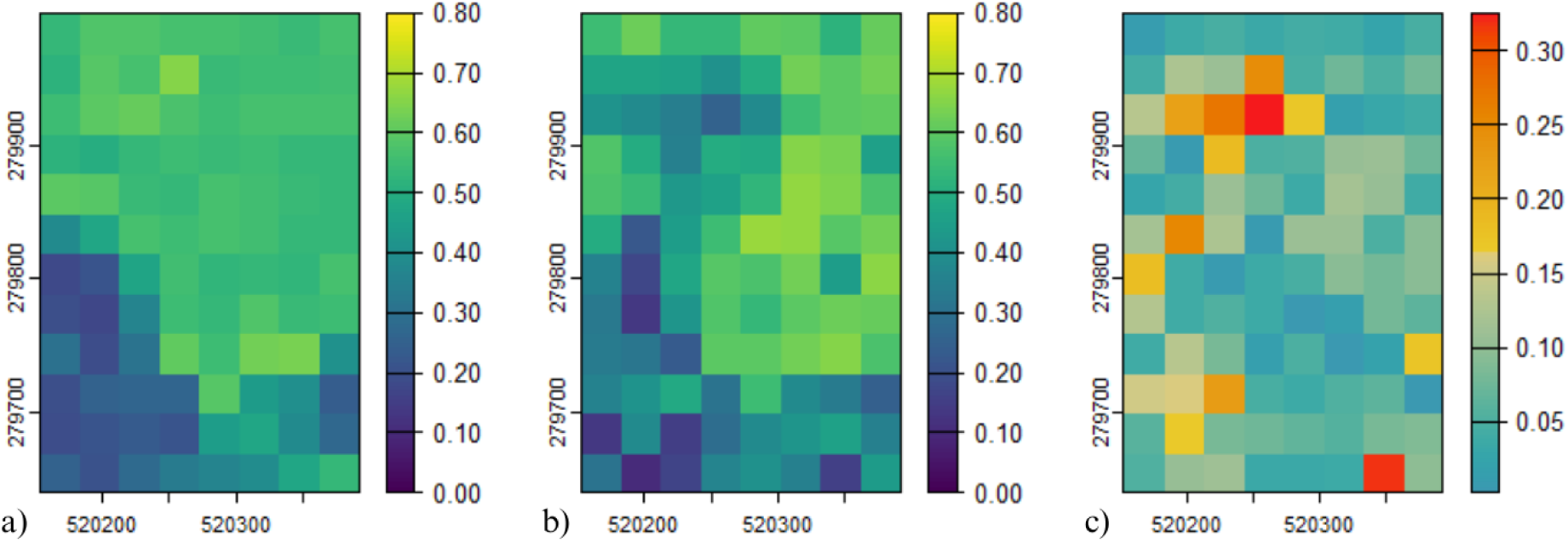
Surfaces depicting the a) predicted canopy cover values from Random Forests for 2015, b) rasterized ALS data for 2015, and c) a difference raster. Pixels are 30 x 30 metres.

**S7 Figure.**
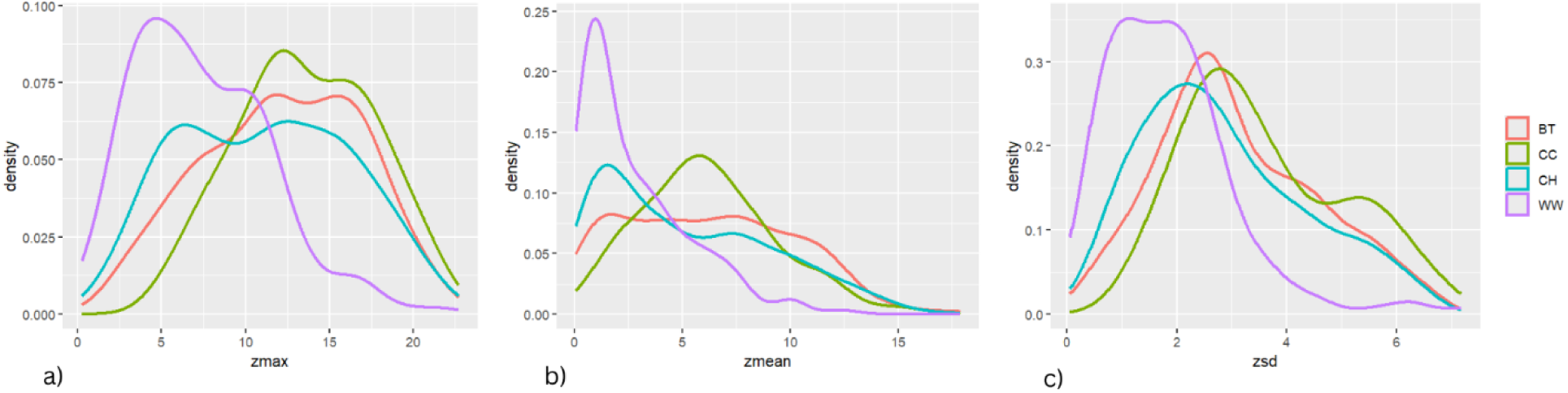
Surfaces depicting the a) predicted canopy relief ratio values from Random Forests for 2015, b) rasterized ALS data for 2015, and c) a difference raster. Pixels are 30 x 30 metres.

**S8 Figure.**
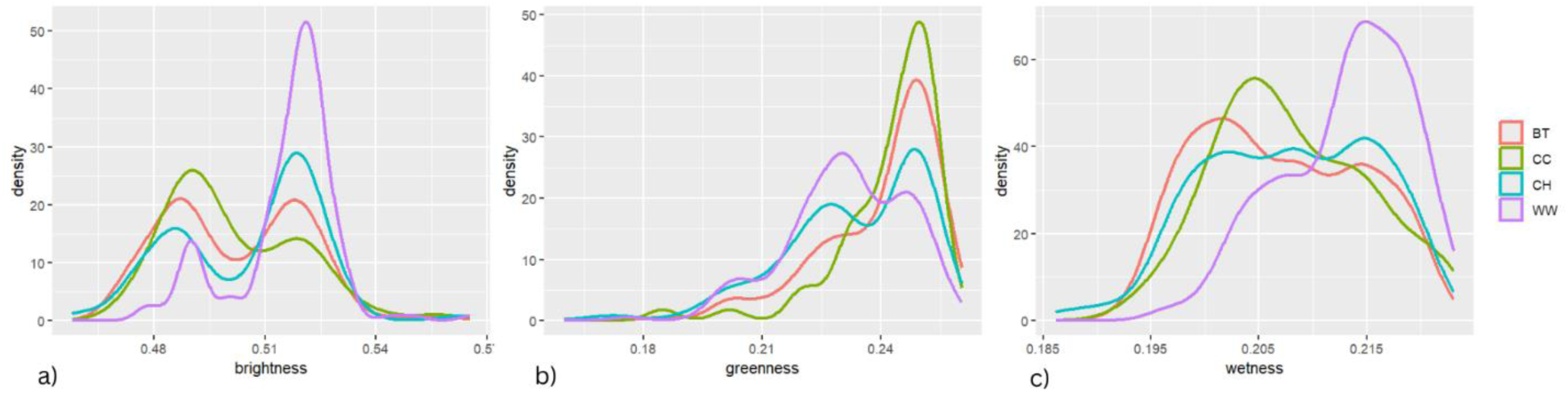
Habitat characteristics for Blue Tit (BT), Chaffinch (CH), Chiffchaff (CC) and Willow Warbler (WW) for shared airborne laser scanning (ALS) important variables: a) maximum height, b) mean height, and c) standard deviation of heights. Y-axis values are density, given that frequencies differ based on number of presences. X-axis values are in metres for the three ALS variables.

**Figure S9.**
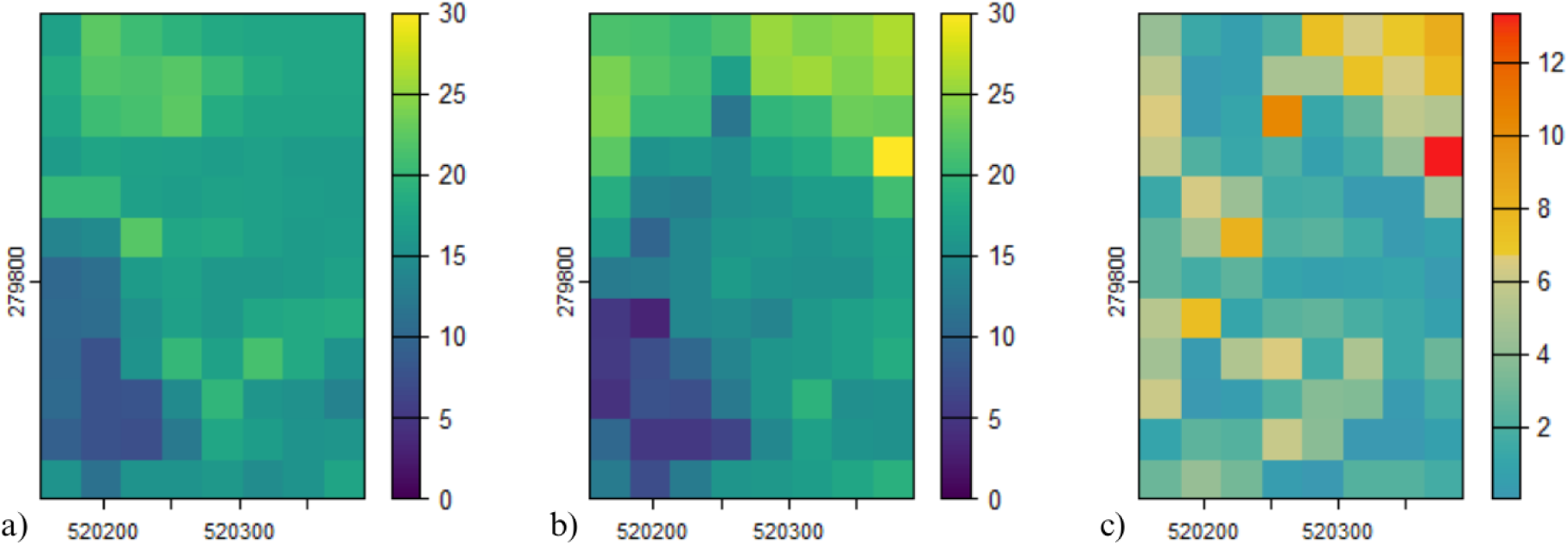
Habitat characteristics for Blue Tit (BT), Chaffinch (CH), Chiffchaff (CC) and Willow Warbler (WW) for shared airborne laser scanning important variables: a) tasseled cap transformation (TCT) brightness, b) TCT greenness, and c) TCT wetness. Y-axis values are density, given that frequencies differ based on number of presences.

